# Epigenetic silencing and blockade of latency reversal by an HIV-1 encoded antisense transcript

**DOI:** 10.1101/2024.12.06.627241

**Authors:** Rui Li, Kaveh Daneshvar, Xinjie Ji, Michelle Pleet, Grace Igbinosun, Mohd. Shameel Iqbal, Fatah Kashanchi, Alan C. Mullen, Fabio Romerio

**Affiliations:** Department of Molecular and Comparative Pathobiology, Johns Hopkins School of Medicine; Baltimore, MD, USA; Division of Gastroenterology, Massachusetts General Hospital; Boston, MA, USA; Department of Biochemistry and Molecular Biology, Johns Hopkins Bloomberg School of Public Health; Baltimore, MD, USA; Laboratory of Molecular Virology, George Mason University; Manassas, VA, USA; Division of Gastroenterology, University of Massachusetts Chan Medical School; Worcester, MA, USA; W. Harry Feinstone Department of Molecular Microbiology and Immunology, Johns Hopkins Bloomberg School of Public Health; Baltimore, MD, USA

**Author notes:** University of Maryland, Baltimore; Baltimore, MD, USA. National Institutes of Health; Bethesda, MD, USA. Summary of the manuscript The HIV antisense transcript AST blocks the reversal of viral latency and can be utilized in ‘block and lock’ cure strategies.

## Abstract

The mechanisms that regulate human immunodeficiency virus 1 (HIV-1) latency are not fully elucidated. We reported that an HIV-1 antisense transcript (AST) induces epigenetic modifications at the HIV-1 promoter causing a closed chromatin state that suppresses viral transcription. Here we show that ectopic expression of AST in CD4+ T-cells from people with HIV-1 under antiretroviral therapy blocks latency reversal in response to pharmacologic and T-cell receptor stimulation, enforcing transcriptional silencing. We define structural domains and sequence motifs of AST contributing to its latency-promoting functions. Finally, we report an unbiased proteomics screen of AST interactors that revealed an array of previously known and potential new HIV-1-suppressive factors. Our studies identify AST as a first-in-class biological molecule capable of enforcing HIV-1 latency and with actionable curative potential.

---

The human immunodeficiency virus type 1 (HIV-1) integrates preferentially in regions of the host genome that contain actively transcribed genes (*1, 2*). At these sites, HIV-1 transcription is regulated through mechanisms that control chromatin organization at the proviral 5’ long terminal repeat (LTR) independently of the surrounding genomic context (*3*). Two nucleosomes, Nuc-0 and Nuc-1 are located at precise positions of the 5’ LTR irrespective of the proviral integration site (*4*). Histone modifications in Nuc-0 and Nuc-1 control the chromatin structure at the HIV-1 promoter and regulate the switch between active transcription and latency (*5, 6*). The ability to modulate the addition or removal of these histone marks can be exploited in HIV-1 cure strategies that seek either to eradicate HIV-1 through latency reversal (*7*) or to force HIV-1 into a state of irreversible latency (*8*). Various classes of latency reversal agents have been evaluated clinically with limited success (*9–12*), whereas strategies aiming at permanent latency via epigenetic silencing of HIV-1 transcription remain largely unexplored.

Trimethylation of histone H3 at lysine 27 (H3K27me3) in Nuc-1 induces a closed chromatin state and suppresses HIV-1 transcription (*13–15*). The Polycomb Repressor Complex 2 (PRC2), which is responsible for depositing the H3K27me3 mark, lacks direct DNA-binding activity (*16*). Long noncoding RNAs (lncRNAs), including natural antisense transcripts (NATs), recruit PRC2 to gene promoters leading to nucleosome assembly, heterochromatinization, and transcriptional silencing (*17, 18*). Studies from our group and others showed that an HIV-encoded antisense transcript (AST) is responsible for the recruitment of PRC2 to the 5’ LTR and for the establishment and maintenance of HIV-1 latency (*19–21*). Here, we present evidence that ectopic expression of AST suppresses HIV-1 transcription, blocking reversal of latency in CD4+ T-cells from people living with HIV-1 (PLWH) under suppressive antiretroviral therapy (ART), and we elucidate the underlying molecular mechanisms.

## Results

### Identification of the AST domain and motifs interacting with the HIV-1 proviral 5’ LTR

We reported that the HIV-1 antisense transcript AST (2574 nt) promotes the establishment and inhibits the reversal of viral latency in *in-vitro* cell line models (*19*). To identify functional determinants of AST, we divided its sequence into five domains: U3 (residues 1-376), A (377-926), B (927-1476), C (1477-2026), and D (residues 2027-2574) (Fig. S1). Secondary structure (minimum free energy) of the mutants was predicted with RNAfold (Fig. S2) (*22–25*). We then generated a battery of deletion and substitution mutants targeting each of the five domains or motifs therein for testing in functional and molecular analyses (Fig. S1 and Table S1). The AST deletion and substitution mutants were then stably transduced into the latently infected cell line, Jurkat E4 (JE4), in which GFP expression serves as readout of viral expression (Fig. S3) (*13*). We have previously used this cell model to demonstrate that AST promotes HIV-1 latency via interaction with PRC2 (*19*).

LncRNAs recruit epigenetic and transcription factors to the chromatin via sequence homology and base pairing with specific genomic DNA regions (*26–29*). The 376-nucleotide (nt) segment at the 5’ terminus of AST that covers part of the U3 region in the 3’ LTR shares perfect sequence identity with the same region of the 5’ LTR, and thus it may be essential for the AST-5’ LTR interaction (Fig. S4). We generated a cell line in the JE4 background stably expressing only the 5’ terminus segment of AST (JE4-U3AST; Fig. S1 and S2) (*19*). This cell line showed a markedly higher percentage of GFP+ cells compared to both parental JE4 cells and JE4 expressing full-length AST (JE4-AST) (Fig. 1A). Higher percentage of GFP+ cells in the JE4-U3AST compared to the other two cell lines reflects the ability of U3AST to interact with the 5’ LTR, displacing the endogenous and biologically active AST that is naturally expressed by the HIV-1 molecular clone in JE4 cells (Fig. S3) (*19*). and the inability of U3AST to recruit silencing factors to the 5’ LTR. Thus, U3AST acts as a dominant negative competitor of endogenous AST, ultimately causing latency reversal.

**Fig. 1.**
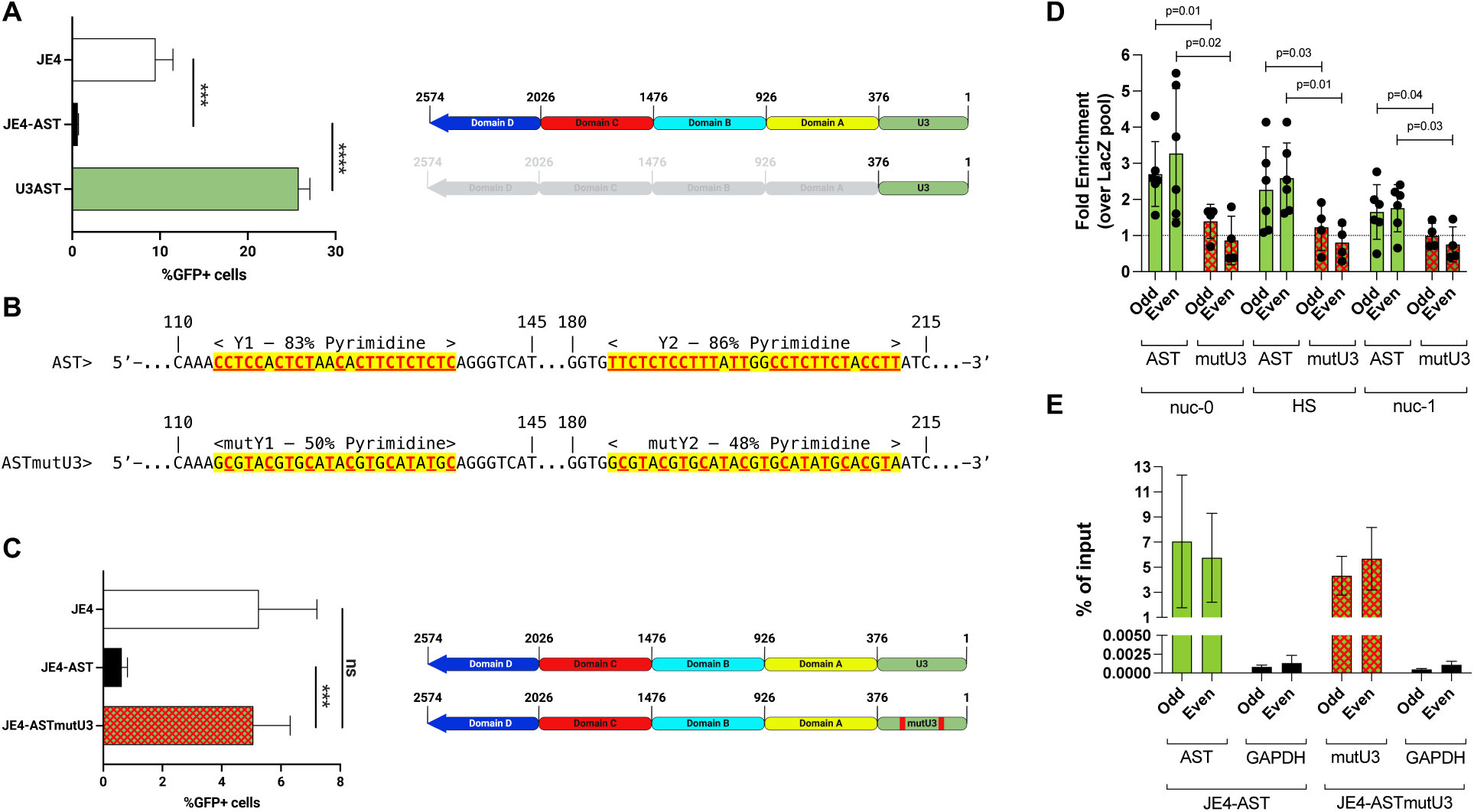
The 5’ terminus domain of AST is required for interaction with the proviral 5’ LTR. (**A**) Cultures of Jurkat E4 cells stably transduced with a lentiviral vector driving ectopic overexpression of AST (JE4-AST) show a lower frequency of productively infected cells compared to parental JE4 cells (∼0.5-1% vs. ∼4-6% GFP+ cells). Cultures of JE4 cells stably transduced with a lentiviral vector expressing only the 376-nt U3 domain at the 5’ terminus of AST (JE4-U3AST) contain a significantly higher percentage of productively infected cells (∼26-28% GFP+ cells). (**B**) Top: nucleotide sequence and position of two pyrimidine rich motifs (Y1 and Y2) in the U3 domain of wildtype AST, which have 83% and 86% pyrimidine content, respectively. Bottom: nucleotide substitutions disrupting the pyrimidine-rich motifs of Y1 and Y2, reducing pyrimidine content in the mutY1 and mutY2 sequences of the ASTmutU3 construct (50% and 48% pyrimidine content, respectively). All the other domains and motifs of ASTmutU3 are the same as in the wildtype AST sequence. Pyrimidines are indicated in red, bold, underlined letters. (**C**) Cultures of JE4 cells stably transduced with a lentiviral vector expressing the ASTmutU3 mutant contain the same percentage of productively infected cells as parental JE4 cells (∼4-6% GFP+ cells). (**D**) ChIRP-qPCR assays show physical interaction of wildtype AST with the Nuc-0 and HS domains in the U3 region of the HIV-1 5’ LTR. Weak interaction between AST and the Nuc-1 domain of the 5’ LTR may reflect incomplete chromatin fragmentation. No interaction is observed between these two domains and ASTmutU3 carrying mutY1 and mutY2 mutations. Results are shown as fold enrichment over LacZ-specific biotinylated antisense oligonucleotide pools. (**E**) RT-qPCR shows equal enrichment of wildtype AST and ASTmutU3 using Odd and Even biotinylated antisense oligonucleotide pools specific for the AST sequence covering domains A through D. No enrichment of GAPDH RNA is observed with either pool of oligonucleotides. ***, p<0.0005; ****, p<0.0001; ns, not significative.

RNA-DNA interactions occur preferentially between pyrimidine-rich (Y-rich) regions in the RNA molecule and purine-rich (R-rich) regions in the DNA (*30–33*). The U3 domain of AST contains two Y-rich motifs (Fig. 1B and S5): a 24-nt Y1 motif (residues 114-137) with 83% pyrimidine content, and a 29-nt Y2 motif (residues 184-212) with 86% pyrimidine content. We generated an AST mutant in which we disrupted the Y1 and Y2 motifs with alternating purine-pyrimidine stretches of equal length (50% and 48% pyrimidine content, respectively; ASTmutU3; Fig. 1B). All the other domains, motifs, and sequences of AST were left unchanged. A JE4-derived cell line stably expressing ASTmutU3 showed a percentage of GFP+ cells higher than JE4-AST and comparable to the parental JE4 cell line (Fig. 1C), suggesting that the disruption of the two Y-rich motifs abolishes the ability of AST to bind the 5’ LTR, reversing the percentage of latently infected cells to baseline levels of the parental JE4 cell line. This conclusion was verified via chromatin isolation by RNA precipitation (ChIRP)-qPCR assays. We generated 24 biotinylated oligonucleotide probes targeting the portion of AST sequence comprised between residues 377 to 2574, which are shared in wildtype AST and ASTmutU3 (Fig. S1, Table S2). The probes were then assigned to either of two pools (Odd or Even), which were used to pull down AST and ASTmutU3 from the respective whole cell lysates. Genomic DNA coprecipitating with the two transcripts was probed by qPCR assays targeting the Nuc-0 (positions –405 to –245 relative to the +1 transcription start site), the DNase I hypersensitivity regions (HS; positions –245 to +1), and the Nuc-1 region (positions +20 to +165) of the 5’ LTR.

We found that AST interacts with the Nuc-0 and the HS sequences in the U3 region of the 5’LTR (∼3-fold and ∼2.5-fold enrichment over LacZ control, respectively; Fig. 1D). Weak interaction with the Nuc-1 region is likely due to incomplete chromatin fragmentation (∼1.5-fold enrichment; Fig. 1D). No binding was observed between ASTmutU3 and the three regions of the 5’ LTR (Fig. 1D). RT-qPCR assays confirmed equal recovery of AST and ASTmutU3 with the two pools of probes, which did not enrich GAPDH RNA negative control (Fig. 1E). Therefore, the U3 domain of AST (residues 1-376) interacts with the proviral 5’ LTR. Two polypyrimidine motifs within this domain are critical for the AST-5’ LTR interaction.

### Identification of the AST domain and motifs interacting with PRC2

Next, we sought to identify the domain of AST that interacts with PRC2. Based on the observation that the JE4 line stably transduced with ASTmutB and to a lesser extent ASTmutA showed significantly higher frequency of GFP+ cells compared to both the parental JE4 and the JE4-AST lines (Fig. 2A), we hypothesized that segment B may be involved in recruiting PRC2 and promoting latency. Replacement of segments C and D with an unrelated exogenous sequence had a much more limited effect (Fig. 2A).

**Fig. 2.**
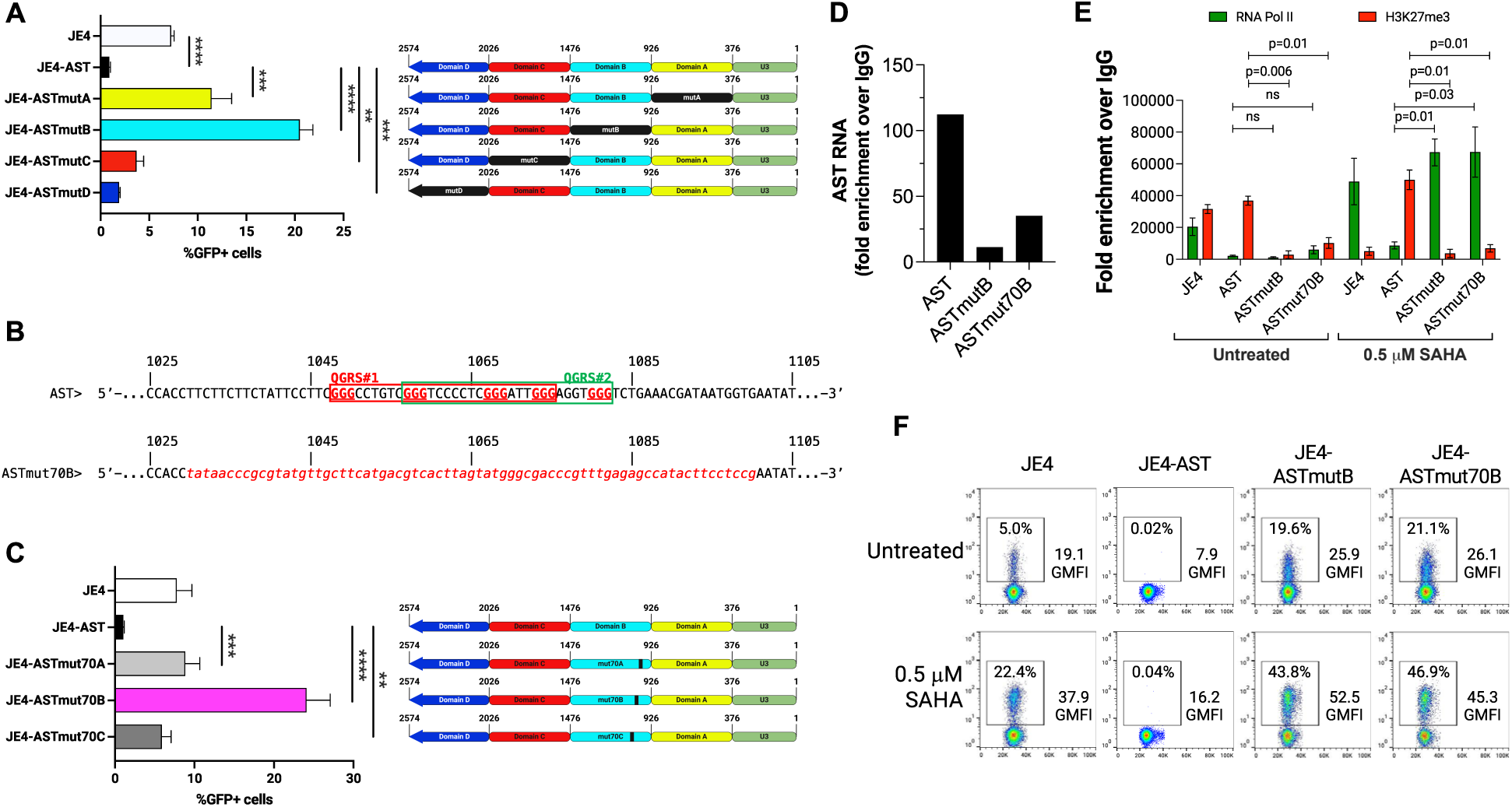
Domain B of AST is primarily responsible for interaction with PRC2. (**A**) Frequency of productively infected (GFP+) cells in cultures of JE4 stably transduced with lentiviral vector expressing AST with substitutions in domains A, B, C, and D (ASTmutA, ASTmutB, ASTmutC, and ASTmutD, respectively) with an exogenous sequence of equal length. Among the four, substitution of domain B (ASTmutB), and to a lesser extent domain A (ASTmutA), shows the most significant impact on AST activity (22-24% and 12-14% GFP+ cells, respectively). Substitution of domains C and D (ASTmutC and ASTmutD, respectively) has a much more limited impact on AST function (∼2-3% GFP+ cells in both cultures). (**B**) Top: nucleotide sequence and position of two overlapping QGRS motifs within domain B of AST (QGRS #1 enclosed in red, and QGRS #2 enclosed in green). G-triplets are indicated in red, bold, underlined letters. Bottom: substitution of the 70-nt sequence encompassing both QGRS motifs in domain B of AST with an exogenous sequence of equal length (ASTmut70B). Sequence substitution is in red italics. All the other domains and motifs of ASTmut70B are the same as in the wildtype AST sequence. (**C**) Cultures of JE4 cells stably transduced with a lentiviral vector expressing the ASTmut70B mutant contain a significantly higher percentage of productively infected cells (∼25-27% GFP+ cells) compared to JE4-AST cells. Substitution of a 70-nt sequence of AST immediately upstream (ASTmut70A) or downstream (ASTmut70B) of the mutation in ASTmut70B shows a much more limited impact (∼2-4% GFP+ cells in both mutants). (**D**) RIP-RT-qPCR assays show physical interaction of wildtype AST with the EZH2 subunit of the PRC2 complex. Such interaction is significantly reduced in cells expressing the ASTmutB or ASTmut70B mutants. (**E**) ChIP-qPCR assays probing H3K27me3 and RNA polymerase II (RNA Pol II) levels at Nuc-1 in JE4, JE4-AST, JE4-ASTmutB, and JE4-ASTmut70B before and after treatment with 0.5 µM SAHA. In JE4 cells, SAHA treatment effectively lowered the levels of H3K27me3 at Nuc-1, whereas in JE4-AST cells they remained high also after SAHA treatment. In the ASTmutB and ASTmut70B mutants, H3K27me3 levels were low both before and after SAHA treatment. Moderate levels of RNA Pol II poised at Nuc-1 were observed in untreated JE4 cells, and they increased after SAHA treatment. RNA Pol II levels at Nuc-1 were low both before and after SAHA treatment in JE4-AST cells. Low levels of RNA Pol II poised at Nuc-1 were observed in untreated JE4 cells, but much higher levels were recruited after SAHA treatment. (**F**) Flow cytometry analyses of GFP expression in JE4, JE4-AST, JE4-ASTmutB, and JE4-ASTmut70B reflect those from ChIP-qPCR assays in (**E**). **, p<0.005; ***, p<0.0005; ****, p<0.0001.

Studies have shown that PRC2 binds G-rich sequences containing [G_3_N_1-5_]≥_4_ motifs able to form G-quadruplex structures (*34–37*). Using the QGRS Mapper server (*38*), we identified two [G_3_N_3-7_]_4_ motifs in AST that match quadruplex forming G-rich sequences (G-score 38 for both) (Fig. 2B and S5). These two motifs are overlapping and map within segment B of AST: one at residues 1048-1075 and the other at residues 1057-1082 (Fig. S5). We replaced a 70-nt sequence of AST encompassing the two overlapping motifs with an exogenous sequence of equal length (ASTmut70B; residues 1031-1100; Fig. 2B, S1 and S2). As controls, we used the same exogenous sequence to replace 70 nt upstream (ASTmut70A; residues 946-1015) or downstream (ASTmut70C; residues 1141-1210) of the mut70B site (Fig. S1 and S2). A JE4-derived cell line stably transduced with the ASTmut70B mutant expressed GFP at significantly higher levels than AST (Fig. 2C). The ASTmut70A and C control mutants had minimal impact (Fig. 2C).

RNA immunoprecipitation (RIP)-RT-qPCR assays with antibodies against EZH2, a core components and catalytic subunit of PRC2 complex, showed significantly reduced enrichment of ASTmutB and ASTmut70B compared to wildtype AST in the EZH2 immunocomplexes (Fig. 2D). Chromatin immunoprecipitation (ChIP)-qPCR assays were performed with cells untreated and treated with the latency reversal agent, suberoylanilide hydroxamic acid (SAHA; 0.5 µM) using antibodies to RNA Polymerase II (Pol II) and H3K27me3 and qPCR primers for Nuc-1. JE4-AST-cells showed high levels of H3K27me3 both before and after SAHA treatment, whereas JE4-ASTmutB and ASTmut70B cells showed the inverse scenario (Fig. 2E). The levels of Pol II at Nuc-1 in JE4-AST cells were very low both before and after SAHA treatment. In untreated JE4-ASTmutB and ASTmut70B cells, RNA Pol II levels at Nuc-1 were low but increased significantly following SAHA treatment (Fig. 2E). Overall, ChIP-qPCR results were consistent with GFP expression levels (Fig. 2F). In particular, high percentage of GFP+ cells in untreated ASTmutB and ASTmut70B cells suggests that the low levels of RNA Pol II observed at Nuc-1 in these cultures (Fig. 2E) is consistent with a fast rate of transcription initiation. The increase in the percentage of GFP+ cells and geometric mean fluorescence intensity (GMFI) after SAHA treatment reflects augmented RNA Pol II recruitment (Fig. 2E). Overall, the two overlapping [G_3_N_3-7_]_4_ motifs in segment B contribute significantly to PRC2 binding and recruitment to the HIV-1 5’ LTR.

### Identification of new AST interactors at the 5’ LTR

Mutation of segments C and D (mutC and mutD) individually as well as deletion of domain D (delD; Fig. S1 and S2) had limited impact on HIV-1 expression (Fig. 2A and 3A). However, mutation or deletion of both domains concurrently (mutCD and delCD; Fig. S1 and S2) had a profound effect (Fig. 3A), suggesting that domains C and D may contribute to the interaction between AST and additional host factors.

**Fig. 3.**
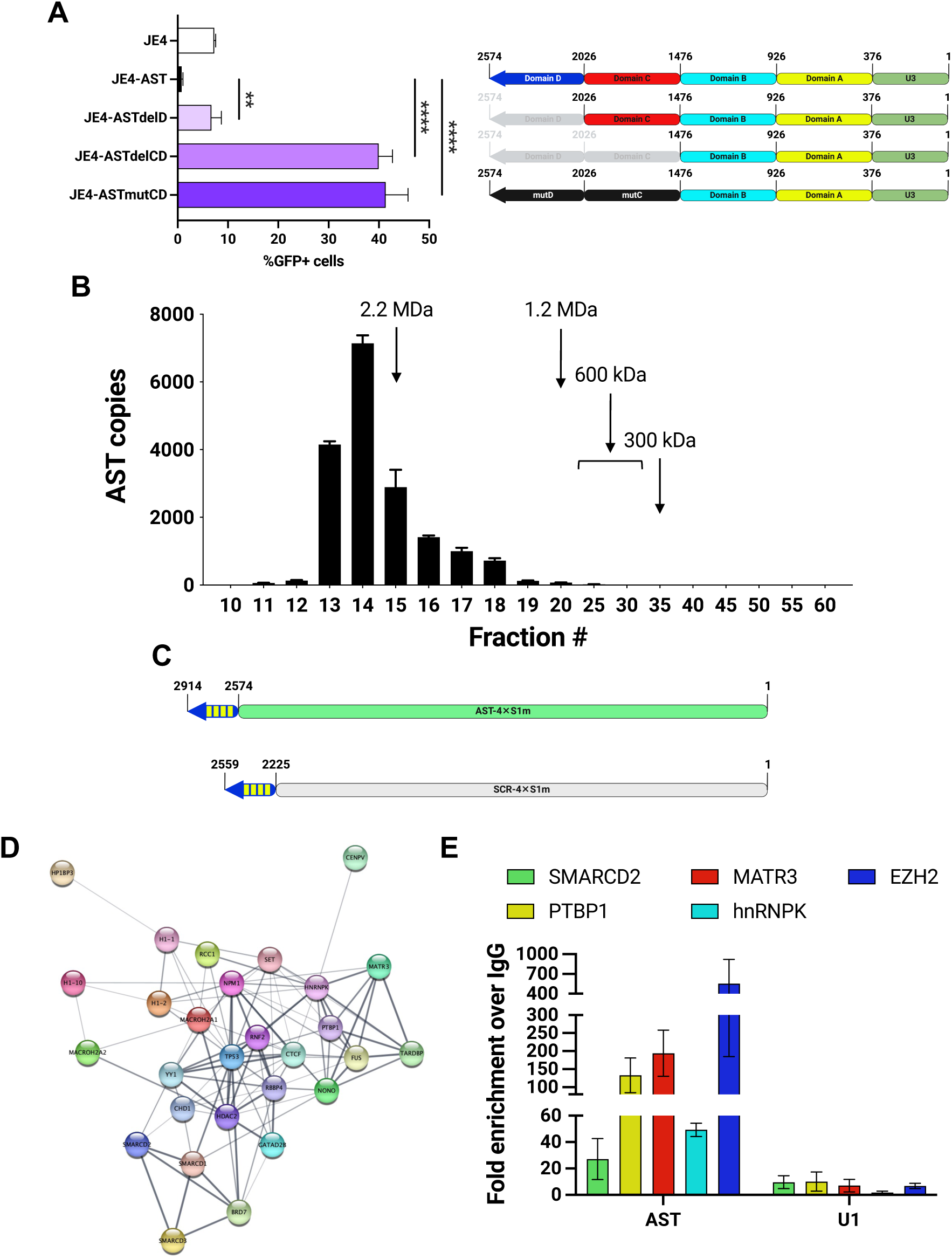
Domains C and D of AST contribute to the recruitment of additional chromatin factors. (**A**) Limited impact of deletion of domain D of AST (ASTdelD) on the frequency of productively infected cells (∼5-7% GFP+ cells). This is consistent with results from ASTmutD cultures. Deletion or substitution of both domains C and D (ASTdelCD and ASTmutCD, respectively) profoundly impacts AST activity (36-38% and 46-48% GFP+ cells, respectively). (**B**) RT-qPCR assays measuring AST levels in size exclusion chromatography fractions from lysates of JE4-AST cells. Greater than 80% of AST molecules elute in fractions 11 through 15, which contain complexes of estimated molecular weight ≥2.2 MDa. (**C**) Schema of AST and SCR molecules with four copies of the streptavidin-binding S1m RNA aptamer (4×S1m) fused at the 3’ terminus. (**D**) STRING network of selected AST interactors included in Table 1. (**E**) RIP-RT-qPCR assays validating interaction between AST and the four new interactors, SMARCD2, PTBP1, MATR3, and hnRNPK selected from the list in Table 1. EZH2 serves as positive control. U1 snRNA is used as a negative control. **, p<0.005; ****, p<0.0001.

**Table 1.** Subset of new AST interactors. New AST-binding factors identified by mass spectrometry in AST pulldown streptavidin complexes. The list of proteins reported here are known to play a role in transcription regulation in various capacities, and it is a subset of AST interactors reported in Data S1. Fold enrichment is relative to SCR-4×S1m control.

| Protein | Fold enrichment | Function |
| --- | --- | --- |
| SMARCD3 | 15.250 | Chromatin remodeling; SWI/SNF complex |
| CENPV | 14.000 | Heterochromatin assembly |
| YY1 | 11.724 | Transcription factor |
| CHD1 | 10.417 | Chromatin remodeling |
| SMARCD1 | 10.339 | Chromatin remodeling; SWI/SNF complex |
| SMARCD2 | 5.370 | Chromatin remodeling; SWI/SNF complex |
| TP53 | 4.974 | Transcription factor |
| CTCF | 4.524 | 3D chromatin regulator |
| HIST1H1A | 4.286 | Histone H1 isoform; higher order chromatin |
| RCC1 | 3.837 | Chromatin condensation |
| H2AFY2 | 3.811 | Histone H2A isoform; transcription repressor |
| GATAD2B | 3.484 | Transcription repressor; NuRD complex |
| RNF2 | 3.465 | Transcription repressor; PcG complex |
| PTBP1 | 3.367 | Gene silencing; biomolecular condensates |
| NONO | 3.535 | Scaffold protein; biomolecular condensates |
| HDAC2 | 3.262 | Histone deacetylase; NuRD complex |
| SET | 3.011 | Gene silencing; biomolecular condensates |
| BRD7 | 2.996 | Transcription factor |
| HP1BP3 | 2.757 | Heterochromatin organization |
| HNRNPK | 2.459 | Gene silencing; biomolecular condensates |
| HIST1H1C | 2.308 | Histone H1 isoform; higher order chromatin |
| MATR3 | 2.308 | Gene silencing; biomolecular condensates |
| NPM1 | 2.207 | Scaffold protein; biomolecular condensates |
| H2AFY | 2.176 | Histone H2A isoform; transcription repressor |
| TARDBP | 2.174 | Transcription repressor; TAR DNA binding |
| FUS | 2.115 | RNA-binding protein; biomolecular condensates |
| RBBP4 | 2.065 | Chromatin remodeling; NuRD, PRC2 complex |
| H1FX | 2.036 | Histone H1 isoform; higher order chromatin |

To address this hypothesis, we fractionated JE4-AST whole cell lysates by size exclusion chromatography. RT-qPCR analyses showed that ∼80% of AST eluted in a very narrow range of fractions containing macromolecular complexes of ≥2.2 million Daltons (MDa) in size (Fig. 3B). Given that the combined molecular weight of AST and the core PRC2 complex is estimated at ∼1MDa (*39*), >1.2MDa remained unaccounted for in the AST-containing complexes that elute in fractions 11-15. This suggests that AST associates with several other host factors in large multimolecular complexes.

To identify new AST interactors, we fused four copies of the streptavidin-binding S1m RNA aptamer at the 3’ terminus of AST and the SCR RNA (AST-4×S1m and SCR-4×S1m; Fig. 3C) (*40, 41*). 4×S1m-containing RNA molecules were pulled down with streptavidin beads from lysates of HEK293T stably transduced with vectors expressing the two chimeric constructs. Co-purifying proteins in the 30-150 kDa range significantly enriched in the AST-4×S1m pull-down were identified by mass spectrometry (MS) (Data S1). These include transcription factors and subunits of epigenetic and chromatin remodeling complexes known to promote HIV-1 silencing and latency, such as components of BAF/PBAF complex (SMARCD1-3), HDAC2, and components of the Polycomb complex (RBBP4 and RNF2) (Fig. 3D and Table 1) (*5, 42–45*). Interestingly, the MS screen also identified additional host factors involved in heterochromatinization of the X chromosome via interaction with the *Xist* lncRNA (*46*): TAR DNA-binding protein 43 (TDP-43), human nuclear ribonuclear protein K (hnRNPK), matrin 3 (MATR3), and polypyrimidine tract binding protein 1 (PTBP1). RIP assays confirmed the interaction between AST and some of these factors (Fig. 3E). These results suggest that the HIV-1-silencing properties of AST may occur via interaction with and recruitment of a macromolecular complex of host factors acting cooperatively.

### Blockade of latency reversal by AST in cells from ART-suppressed PLWH

We reported that the HIV-1 antisense transcript AST (2574 nt) promotes the establishment and inhibits the reversal of viral latency in *in-vitro* cell line models (*19*). Here, we assessed whether AST is able to maintain HIV-1 in a latent state and block reactivation of viral transcription in resting CD4+ T-cells freshly isolated from peripheral blood of PLWH undergoing suppressive ART for at least 1 year (median 6 years; range 1-18 years; Table S3). A vector expressing either AST or a control scrambled sequence (SCR) under the eukaryotic translation initiation factor 1A (EIF1α) promoter was introduced via nucleofection into freshly isolated, resting CD4+ T-cells. Both vectors also express green fluorescent protein (GFP) under the human cytomegalovirus (CMV) promoter (Fig. 4A). Latency reversal was then triggered via treatment with the latency reversal agents SAHA and Panobinostat, or through T-cell receptor (TCR) stimulation with anti-CD3/CD28-coated magnetic beads (Fig. 4A). All three treatments reactivated HIV-1 transcription to various degrees in donor cells expressing SCR, but not AST (Fig. 4B; p<0.002, p=0.016, p=0.031 for SAHA, Panobinostat, and anti-CD3/CD28, respectively). Efficiency of nucleofection was assessed as the percentage of GFP-positive cells, and it ranged between 20-40% (Fig. 4C and S6). Given the expected low levels of protein translation in resting CD4+ T-cells, expression of GFP is likely to underestimate the efficiency of nucleofection. For a more accurate estimate that can account for the near-complete blockade of latency reversal in cells receiving the AST construct, we labeled the two plasmid vectors with the fluorescent dye, Cyanine-5 (Cy5), and we assessed the percentage of Cy5-positive cells after nucleofection (Fig. 4C and S7). The estimate of nucleofection efficiency through this method was consistently between 75-90%, and it was 1.5- to 3-fold greater than the one based on GFP expression (p=0.001 and p=0.0005 for cells transfected with SCR and AST vectors, respectively). Expression levels of both AST and SCR were analyzed by strand-specific RT-qPCR in both cell cultures (Fig. 4D). As previously reported (*19*), we detected endogenous expression of AST in CD4+ T-cells from PLWH after nucleofection with SCR-expressing vector. Much higher expression levels were detected after nucleofection of the AST expression vector (p-0.0003). SCR RNA was readily detected following nucleofection of SCR-expressing vector, but only at background levels in AST-nucleofected cells (p=0.008). Altogether, these results show that ectopic overexpression of AST in cells from ART-suppressed PLWH effectively blocks reversal of latency and viral transcription in response to various stimuli.

**Fig. 4.**
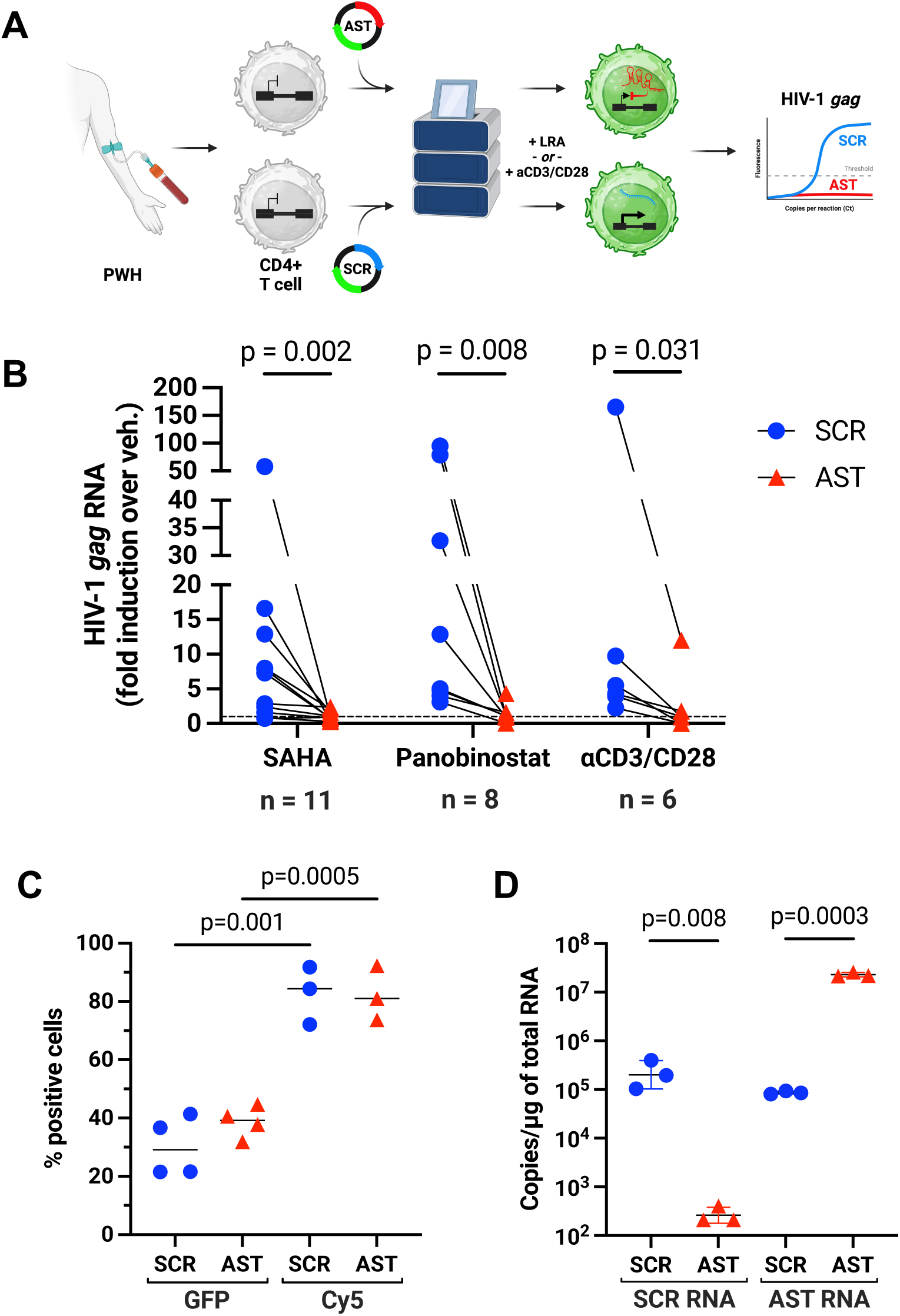
Blockade of latency reversal in CD4+ T-cells from PLWH after ectopic expression of AST. (**A**) Diagram of experimental design. Peripheral blood was collected from 15 people living with HIV-1 (PLWH) under suppressive antiretroviral therapy (Table S3), and CD4+ T-cells were isolated by negative selection. After nucleofection with vectors expressing either AST or SCR, cells were treated with latency reversal agents (LRA; SAHA or Panobinostat) or magnetic bead conjugated anti-CD3/CD28 antibodies. Levels of HIV-1 *gag* RNA were analyzed by semi-nested RT-qPCR. (**B**) HIV-1 *gag* RNA expression in CD4+ T-cells from PWH nucleofected with vectors expressing AST (red triangles) or an unrelated control transcript (SCR, blue circles) and then treated with SAHA (left; *n* = 11), Panobinostat (middle; *n* = 8), or anti-CD3/CD28 antibodies (right; *n* = 6). RNA levels are expressed as fold increase over vehicle control (DMSO for SAHA and Panobinostat, and PBS for anti-CD3/CD28). P-values were determined with the Wilcoxon matched pairs signed rank test. (**C**) Efficiency of nucleofection in freshly isolated, resting CD4+ T-cells from PLWH. This was determined by both expression of GFP from the same vector also expressing AST or SCR (Fig. S6), and by intracellular Cy5 content when using Cy5-labeled vectors in nucleofection (Fig. S7) Frequency of transfected cells measured with the two methods were significantly different both for cells transfected with SCR vector and cells transfected with AST vector (p=0.001 and p=0.0005, respectively). (**D**) Quantitative RT-qPCR assays measuring AST and SCR RNA expression levels in freshly isolated, resting CD4+ T-cells from PLWH nucleofected with SCR- or AST-expression vectors. The levels of the two transcripts were significantly different between cells nucleofected with each of the transcript-expressing vectors and cells nucleofected with the other transcript-expressing vector (p=0.008 for SCR levels in SCR-nucleofected cells vs. AST-nucleofected cells; p=0.0003 for AST levels in AST-nucleofected cells vs. SCR-nucleofected cells). Higher baseline levels of AST in SCR-nucleofected cells reflect endogenous expression of AST in PLWH-derived samples.

## Discussion

Antisense RNA-based mechanisms ensure a tight control of gene expression by establishing a threshold of activation that buffers weak stimuli and represses stochastic expression of the target gene (*47, 48*). Stimuli that exceed such thresholds switch on gene expression, which rapidly reaches maximal levels (*49, 50*). As stimuli subside, antisense transcripts accelerate the return to baseline levels (*51*). *Xist*, *ANRIL*, and *Kcnq1ot1* are examples of antisense transcripts that function in such manner via recruitment of suppressive epigenetic factors that trigger a closed chromatin state and silence transcription (*29, 52–54*).

The impact of AST on HIV-1 transcription involves mechanisms similar to those described for eukaryotic antisense transcripts. AST is expressed constitutively from a Tat-independent negative sense promoter in the 3’ LTR (*19, 20, 55–57*). However, we showed that AST levels increase significantly in response to several stimuli (*58*), similarly to other antisense transcripts with gene regulatory function (*29*). AST has a predominantly nuclear localization due to inefficient polyadenylation (*59*) and, possibly, interaction with nuclear RNA-binding proteins (Table 1 and Data S1) (*60*). Our previous studies using cell line models demonstrated that AST promotes HIV-1 latency by recruiting the chromatin remodeling complex, PRC2 to the 5’ LTR (*19*). Here we provided proof that AST can also block latency reversal in an *ex-vivo* model based on CD4+ T-cells from ART-suppressed PLWH. This ability is especially remarkable in consideration of the HIV-1 sequence variability within and among infected individuals.

Owing to their complex secondary and tertiary structure and their length, lncRNAs serve as scaffolds for the recruitment of proteins that regulate multiple biological functions, including DNA repair and replication, transcription, and chromatin remodeling (*61, 62*). Above a threshold concentration, lncRNAs can selectively partition specific proteins into biomolecular condensates that are formed by phase separation (*63, 64*). By contrast, knocking down expression of lncRNAs or deleting essential protein-interacting domains prevents formation of the condensates and their function (*65–67*). The lncRNAs *eRNA* (*67*), *HSATII* (*68*), *HOTAIR* (*69*), *NEAT1* (*70*), *DIGIT* (*41*), and *Xist* (*71*) regulate transcription through their interaction with RNA binding proteins (RBP) that contribute to the recruitment of chromatin remodeling factors (*29, 61*).

Our studies suggest that AST may promote the formation of biomolecular condensates that negatively regulate the 5’ LTR promoter activity. We showed that AST is associated exclusively with large macromolecular complexes of >2MDa. In addition, we found that AST interacts with RBPs such as hnRNPK, MATR3, PTBP1, TDP-43, FUS, NONO, and NPM1 (Table 1 and Data S1), known to cooperate with lncRNA during phase separation and formation of biomolecular condensates (*29, 61, 62*). Deletion or mutation of structural domains and sequence motifs of AST eliminates its ability to promote latency. In a previous study we also identified and mapped AST epigenetic and post-transcriptional modifications, including 2′-*O*-ribose methylation (Nm), adenosine-to-inosine (A-to-I), and pseudouridine (Ψ) (*72*), which are catalyzed by enzymes identified in the screen of AST interactors (FBL, FTSJ3, and DKC1; Data S1) and that have been shown to facilitate lncRNA-RBP interactions (*73–81*).

Cure and treatment strategies that seek to permanently suppress viral transcription remain underdeveloped compared to those proposing to flush out or disable proviruses hiding in cellular and anatomical reservoirs (*7, 8, 82, 83*). This is especially true for approaches that aim to achieve epigenetic silencing of the HIV-1 promoter (*84, 85*). Our published studies and the ones described here show that AST establishes a tight control of HIV-1 transcription, locking the provirus in a latent state refractory to external stimuli and accelerating its entry into latency when viral expression wanes (*19*). Ectopic expression of AST at high levels is expected to augment the local concentration of epigenetic repressors at the 5’ LTR and to block transcription. Altogether, this provides a rationale for the use of AST as a curative agent that can lock HIV-1 in a state of permanent latency.

## Author contributions

Conceptualization: FR, RL

Methodology: RL, KD, MP

Investigation: RL, KD, MP, XJ, GI, MSI

Visualization: RL, FR

Funding acquisition: FR, FK, ACM

Project administration: FR

Supervision: FR, FK, ACM

Writing – original draft: FR

Writing – review & editing: All authors

Competing interests

Authors declare that they have no competing interests.

Data and materials availability

All data are available in the main text or the supplementary materials.

Supplementary Materials

## Materials and Methods

Figs. S1 to S7

Tables S1 to S5

Data S1

References: 86-88

## Materials and Methods

### Participant cohorts, sample collection, and study approval

Peripheral blood was collected from PLWH on ART, who were continually suppressed (<50 copies HIV-1 RNA copies per mL) for at least 1 year at the time of blood collection (Table S3). Donors were enrolled at the Johns Hopkins Hospital after approval from the Johns Hopkins Institutional Review Board (protocol # IRB00280013). All donors provided written informed consent.

### Isolation of resting CD4+ T cells, nucleofection, latency reversal, and HIV-1 RT-qPCR

PBMC were isolated from peripheral blood by Ficoll centrifugation and used to enrich resting CD4+ T cells by negative selection via MACS (Miltenyi). Untouched CD4+ T cells were then nucleofected with a vector expressing AST (*19*) or SCR under the EF1α promoter using the Amaxa P3 Primary Cell nucleofection kit and an Amaxa Nucleofector 4D (Lonza) programmed for an EH-100 pulse code. After 48 hours, cells were treated with 0.5 µM SAHA (BEI Resources, NIH HIV Reagent Program, Division of AIDS, NIAID, NIH: ARP-12130), 30 nM Panobinostat (Selleckchem), and anti-CD3/CD28 Dynabeads (1 bead/cell; Thermo Fisher). As vehicle control for cells treated with SAHA and Panobinostat, cells were treated with DMSO (1:2000; Sigma Aldrich). As vehicle control for cells treated with anti-CD3/CD28 and Panobinostat, cells were treated with an equal volume of PBS (Thermo Fisher). All treatments were performed in the presence of 2 µM raltegravir (BEI Resources, NIH HIV Reagent Program, NIAID, NIH: HRP-11680) and 1 U/ml IL-2 (Thermo Fisher). Nucleofected cells treated with vehicle were used to determine baseline HIV-1 transcription. After 72 hours, cells were collected in Trizol (Thermo Fisher Scientific) and total RNA was extracted using the Dircted-Zol RNA Microprep kit (Zymo Research) with on-column DNase treatment. Fifty nanograms of total RNA were reverse transcribed into cDNA using Superscript III Reverse Transcriptase (Thermo Fisher Scientific). Briefly, RNA was incubated at 65°C for 5 min with 1.5 µg random hexamers, 0.25 µg Oligo dT(12–18) before adding first strand buffer, 5 mM DTT, 20 U RNAseOUT RNAse inhibitor and 5% RT enzyme in a volume of 20 µL. The reaction was incubated at 42°C for 45 min and 80°C for 5 min. The cDNA was then used to perform a semi-nested real time quantitative qPCR assay for cell-associated unspliced HIV-1 Gag RNA as previously described (*11, 86*) with primers in Table S4. The PCR conditions for Gag RNA were: 10% Amplitaq Buffer II, 2.5 mM MgCl_2_, 0.2 mM of each dNTP, 0.4 µM of each primer and 0.5% Amplitaq Gold polymerase in a reaction volume of 50 µL and run at 95°C for 10 min, followed by 15 cycles of 94°C for 20 seconds, 55°C for 40 seconds, and 72°C for 40 seconds. For the second round PCR, 2 µL of first round PCR product was used as DNA template with 47% PowerUp Sybr Green master mix (Thermo Fisher Scientific) and 0.75 µM of each primer. qPCR settings were as follows: 50°C for 2 minutes, then 95°C for 10 minutes, followed by 40 cycles of 95°C for 20 seconds, 60°C for 40 seconds. All samples were run together with no RT control wells to assess for DNA contamination. Expression data for the fold change over vehicle control was calculated using 2^-ΔΔCt^ method.

### Cell lines, reagents, lentiviral transduction, flow cytometry

Lenti-X 293T (Takara) were cultured in DMEM supplemented with 10% fetal bovine serum (FBS), penicillin, streptomycin, and L-glutamine. The latently infected cell line Jurkat E4 (a kind gift of Jonathan Karn, Case Western Reserve University, (*6, 13*)) were cultured in RPMI 1640 supplemented with 10% FBS and antibiotics. Lentiviral transduction was carried out as we described previously (*19*). Briefly, AST-derived sequences were cloned into the pLVX-Puro vector (Takara) and then transfected into Lenti-X 293T cells using the Lenti-X Packaging Single Shots (VSV-G) system (Takara). Culture supernatants were collected, and viral particles concentrated using the PEG-*it* reagent (System Biosciences). Jurkat E4 cells were then transduced with purified lentiviral particles overnight in complete RPMI medium containing 2 µg/ml polybrene. Stably transduced cells were selected and maintained in medium containing 1 µg/ml puromycin.

### Flow cytometry

Plasmid DNA was labeled with DNA Label IT Tracker Intracellular Nucleic Acid Localization kit (Mirus, Madison, WI) using 0.5 µl Cy5/µg of plasmid DNA according to the manufacture’s protocol. Unreacted Label IT Tracker reagent was removed by ethanol precipitation. For flow cytometry analyses, samples were acquired on a LSRII Fortessa flow cytometer (BD Biosciences, San Jose, CA). Doublet cells were excluded using forward scatter and side scatter. DNA uptake into cells was determined by analysis of APC positive cells. Raw data were analyzed with FlowJo software.

### RNA isolation, reverse transcription, and quantitative PCR

RNA isolation and RT-qPCR were carried out as described (*19*). Briefly, RNA was isolated using the RNeasy Mini Kit (Qiagen), digested with Turbo DNase (Ambion), and further purified with the RNase Mini Kit. 100-500 ng of RNA were utilized for reverse transcription using the iScript Select cDNA Synthesis Kit (BioRad) and the AST-tag RT primer. For the TaqMan PCR reaction, we utilized the TaqMan Gene Expression Master Mix (Applied Biosystems) according to the manufacturer’s instructions. For quantitative PCR, the AST amplicon carrying the tag sequence was cloned into pUC57. The plasmid was then linearized and utilized at 10^0^-10^5^ copies to generate a PCR standard curve. The copy number per 3 µl of cDNA was then normalized to the equivalent number of cells, which were measured manually with a hemacytometer. For RT-qPCR of the U1 snRNA, RT reaction was carried out with the same kit as above, and with the random primers contained in the kit. The sequence of primers and probe are reported in Table S4. Quantitative TaqMan PCR was carried out as described above.

### RNA immunoprecipitation (RIP)

Five million cells were washed twice with ice-cold PBS and lysed with a buffer containing 100 mM KCl, 5mM MgCl_2_, 10 mM Hepes pH 7, 0.5% Igepal CA-630, 1mM DTT, Protease Inhibitors, and RNaseOUT in molecular biology grade water. After 5 minutes incubation on ice, the lysate was frozen at -80°C overnight. Fifty µl Protein A+G Magnetic Beads were washed with 500 µl of NT-2 buffer (50 mM Tris-HCl pH7.4, 150 mM NaCl, 1mM MgCl_2_, and 0.05% IGEPAL CA-630 in molecular biology grade water) and incubated with 5 µg antibody of pre-immune rabbit IgG or target-specific antibodies in Table S5 with rotation at room temperature for 1 hour. Excess antibody was removed with six washes with NT-2 buffer on ice. After the last wash, 900 µl NET-2 buffer (20 mM EDTA pH8, 1 mM DTT, and RNaseOUT in NT-2 buffer) was added to the antibody-dynabeads complex. The lysate was thawed at room temperature and centrifuged at 14,000 rpm for 10 minutes at 4°C. One hundred µl supernatant was added to the antibody-dynabeads complex followed by overnight incubation with rotation at 4°C and 10 µl supernatant was saved at -80°C as input control. After washing six times on ice, the protein-antibody-dynabeads complex and input were digested with 150 µl proteinase K buffer (1% SDS and 180 µg Proteinase K solution in NT-2 buffer) at 55°C for 30 minutes. The supernatants were transferred to new Eppendorf tubes and vortexed with 250 µl NT-2 Buffer and 400 µl Phenol/Chloroform 5:1 (Fisher BioReagents) for 15 seconds, followed by a 10-minute centrifugation at 14,000 rpm at room temperature. Three hundred and fifty µl of the aqueous phase was collected and vortexed with 400 µl chloroform for 15 seconds, followed by another 10-minute centrifugation at 14,000 rpm at room temperature. Three hundred µl of the aqueous phase was collected for an overnight RNA precipitation at -80°C with 50 µl of 5 M ammonium acetate, 15 µl of 7.5 M LiCl, 5 µl of 1 mg/ml glycogen, and 850 µl of ethanol. RNA was pelleted down by centrifuging at 14,000 rpm for 30 minutes at 4°C and washed with 500 µl 80% ethanol. The RNA pellet was centrifuged again at 14,000 rpm for 15 min at 4°C to remove the supernatant and left on the bench to air dry. Finally, RNA was resuspended in 15 µl molecular biology-grade water for analysis.

### Chromatin immunoprecipitation (ChIP)

Twenty million cells were placed in two 150-mm culture dishes in 20 ml medium and 540 µl of 37% formaldehyde (Sigma, cat. 252549) were added to each culture dish and incubated for 10 minutes at room temperature with gentle shaking. Two ml of 10× glycine (Cell Signaling, 7005) were added to each culture dish and incubated for 5 minutes at room temperature with gentle shaking. Crosslinked cells were combined and pelleted at 1,000×*g* for 5 minutes at 4°C and washed twice with ice-cold PBS. Cell pellet was resuspended in 1 ml of 1× ChIP Sonication Cell Lysis Buffer (Cell Signaling, 96529) with Protease Inhibitor Cocktail (Cell Signaling, 7012) and stored at –80°C if not processed immediately. Cell suspension was incubated on ice for 10 minutes and then pelleted down at 5,000×*g* for 5 minutes at 4°C. The pellet was resuspended in 1 ml of 1× ChIP Sonication Cell Lysis Buffer with PIC and incubated on ice for another 10 minutes. The cell suspension was centrifuged at 5,000×*g* for 5 minutes at 4°C. The pellet was resuspended in 1 ml of ice-cold 1× ChIP Sonication Nuclear Lysis Buffer (Cell Signaling, 28778) with PIC, and aliquoted into four 500 μl PCR tubes for sonication. Lysates were sonicated at 70% amplitude with 15 seconds on and 30 seconds off cycles for 25 minutes. After sonication, lysates were combined into a 1.5 ml Eppendorf tube and clarified by centrifugation at 21,100×*g* for 10 minutes at 4°C. Supernatant was collected to a new tube and stored at –80°C if not processed immediately. Fifty µl of the clarified lysate was mixed with 100 µl of nuclease-free water, 6 µl of 5 M NaCl (Cell Signaling, 7010), and 2 µl of RNase A (Cell Signaling, 7013) and incubated at 37°C for 30 minutes. Then, 2 µl of Proteinase K (Cell Signaling, 10012) were added and sample was incubated at 65°C for 2 hours to overnight. After incubation, chromatin was purified with QIAquick PCR Purification Kit (Qiagen, 28106). Fifteen µl of purified chromatin was taken to determine DNA fragment size by electrophoresis on a 1% agarose gel. Chromatin immunoprecipitation was performed if 60-90% of total DNA fragments are <1 kb. For chromatin immunoprecipitation, sonicated chromatin was diluted with 1× ChIP Buffer (Cell Signaling, 7008) at a dilution factor of 1:4. Ten µl of diluted chromatin were transferred to a microcentrifuge tube as 2% Input and stored at – 80°C until future use. Five hundred µl of diluted chromatin were transferred to microcentrifuge tubes with immunoprecipitating antibodies (Table S5) and incubated at 4°C with rotation for 4 hours to overnight. Then, 30 µl of Protein G Magnetic Beads (Cell Signaling, 9006) were added to each IP reaction and incubated 4°C with rotation for 2 hours. Beads were recovered with a magnetic separation rack and washed three times with low salt wash (1 ml of 1× ChIP Buffer) and once with high salt wash (1 ml of 1× ChIP Buffer plus 70 µl of 5 M NaCl). One hundred fifty µl of 1× ChIP Elution Buffer (Cell Signaling, 7009) was added to each IP sample and the 2% input sample. To elute chromatin from the antibody/Protein G Magnetic Beads, samples were incubated for 30 minutes at 65°C with gentle vortexing (1200 rpm). Supernatant was collected and reverse crosslinking was performed to all samples including the 2% input by adding 6 µl of 5M NaCl and 2 µl Proteinase K followed by a 2 hour to overnight incubation at 65°C. DNA samples were purified with QIAquick PCR Purification Kit and used for qPCR.

### Chromatin isolation by RNA precipitation (ChIRP)

ChIRP assays were carried out using the Magna ChIRP™ RNA Interactome Kit (Millipore Sigma) following the manufacturer’s instructions. Briefly, 2×10^7^ cells were washed with PBS, resuspended in 1% glutaraldehyde in PBS, and incubated at room temperature for 10 minutes. The crosslinking agent was blocked by the addition of glycine, and the cells were washed with PBS. After resuspension in PBS, the cells were lysed, and genomic DNA sheared by sonication with a Q800R Sonicator (Qsonica). Cell lysates were then incubated with Odd and Even pools of AST-specific biotinylated probes (ChIRP Probe Designer, www.biosearchtech.com; Table S2) or LacZ-specific biotinylated probes as a negative control (provided in the kit). After 4 hours hybridization at 37°C with rotation, streptavidin magnetic beads were added, and the hybridization mixture was incubated for an additional 30 minutes at 37°C. The beads and captured material were collected with a magnet and then washed four times. After resuspension in wash buffer, 90% of beads were used for DNA extraction using reagents provided in the ChIRP kit and the remaining 10% for RNA extraction using a miRNeasy® Mini Kit (Qiagen) and. Finally, DNA was analyzed by qPCR using primers specific for Nuc-0, HS, and Nuc-1 (Table S4). Results were expressed as enrichment with AST-specific probes over LacZ control pools. RNA was analyzed by RT-qPCR with primers specific for AST and GAPDH (Table S4) to evaluate efficiency and specificity of RNA recovery.

### Fractionation of whole cell lysates by size exclusion chromatography

JE4-AST cells were washed 3 times with PBS without Mg^2+^ and Ca^2+^ and lysed in a buffer containing 50 mM Tris– HCl pH 7.5, 120 mM NaCl, 5 mM EDTA, 0.5% NP-40, 50 mM NaF, 0.2 mM Na_3_VO_4_, 1 mM DTT, and one complete protease cocktail tablet per 50 ml. After incubation on ice for 20 minutes with gentle vortexing, cell lysates were cleared by centrifugation at 10,000 rpm for 10 min at 4°C. Supernatants were transferred to a clean tube, and protein concentration was determined by Bradford assay (Bio-Rad, Hercules, CA, USA). For size-exclusion fractionation, 2.5-5.0 mg of total protein were diluted to 1 ml total volume with chromatography running buffer (0.2 M Tris– HCl pH 7.5, 0.5 M NaCl, and 5% glycerol) and run on a Superose 6 10/300 size-exclusion chromatography column (GE Healthcare Bio-Sciences, Uppsala, Sweden) using the ÄKTA Purifier system (GE Healthcare Bio-Sciences, Piscataway, NJ, USA). A quarter-inch gap was introduced to the top of the Superose 6 column to better separate small molecular weight complexes from fractions eluting off the far-right side of the chromatogram. After sample injection (using a 1-ml loop), running buffer was set at a flow rate of 0.3 ml/min, and 0.5-ml fractions of the flow-through were collected at 4°C for a total of approximately 60 fractions. Total RNA was extracted with a RNeasy Mini Kit (Qiagen) from 100 µl of every fifth chromatography fraction, and AST copies were measured in 5 µl of purified RNA by RT-qPCR. Given that AST copies peaked at fraction #15, RNA extraction and RT-qPCR were repeated with chromatography fractions 11 to 19.

### Identification of AST-binding partners by 4×S1m-mediated enrichment and mass spectrometry (MS)

The AST-4×S1m and SCR-4×S1m constructs were cloned into the pLVX-Puro vector and stably transduced into Lenti-X 293T cells as described above. For RNA pull-down and mass spectrometry analyses, cells were plated on ten 10-cm^2^ plates and allowed to grow to near confluency. Cells were washed with DPBS and irradiated with 400 mJ/cm^2^ of energy on ice in an ultraviolet cross-linker (Stratalinker). Nuclei of ultraviolet cross-linked cells were isolated by suspending the cells in a hypotonic buffer (20 mM Tris-HCl pH 7.4, 10 mM NaCl and 3 mM MgCl_2_) for 15 minutes, adding NP-40 to a final concentration of 0.5% and then centrifuging for 10 min at 850×*g*. Nuclei were lysed in lysis buffer (150 mM KCl, 25 mM Tris-HCl pH 7.4, 5 mM EDTA, 5 mM MgCl_2_, 1% NP-40, 0.5 mM dithiothreitol (DTT), Roche mini-tablet protease inhibitor and 100 U/ml RNaseOUT) and debris was cleared by centrifugation at 16,000×*g*. The supernatant containing the nuclear lysate was then cleared with 50 µl of Avidin Agarose beads (Thermo Fisher Scientific, 20219) to deplete biotin from the lysates (*87*). The cleared lysate was incubated with 150 µl of streptavidin C Dynabeads (Thermo Fisher Scientific) for 4 hours on a rocker at 4°C. The beads were collected using a magnet and washed three times with a wash buffer of the same composition as the lysis buffer, except KCl was increased to 350 mM. To release the proteins, beads were incubated with RNase A and resuspended in 2× LDS sample buffer. The entire lysates were resolved on a polyacrylamide gel. Segments of the gel from 30 kDa to 150 kDa were excised and analyzed using MS, which was performed at Harvard Medical School Taplin Mass Spectrometry Core. Analysis of the proteomic data was performed using the CRAPome proteomic analysis (*88*), using a threshold of >90% confidence and greater than fourfold enrichment over controls.

### Generation of AST mutants

All AST mutants were generated by DNA synthesis (GeneScript), cloned into the lentiviral vector pLVX-Puro (Takara) under the EIF1α promoter, and stably transduced into JE4 cells as previously described (*19*). All experiments were carried out with the bulk population of stably transduced cells obtained under puromycin selection without isolation of single cell-derived clones. The complete sequence of the wildtype AST and of each AST mutant is reported in Table S1. To generate the ASTmutA, ASTmutB, ASTmutC, and ASTmutD mutants we replaced each domain with the same exogenous sequence of 550 nt. To generate the ASTmutCD mutant, domains C and D were replaced with two copies of the 550-nt exogenous sequence placed in tandem. To generate the ASTmut70A, ASTmut70B, and ASTmut70C mutants, we used the first 70 nt of the exogenous sequence above. The ASTmutU3 mutant was generated by replacing the Y1 (residues 114-137 of the full-length AST; 83% pyrimidine content) and Y2 motifs (residues 184-212 of the full-length AST; 86% pyrimidine content) with sequences of alternating purine-pyrimidine residues (50% and 48% pyrimidine content, respectively) of equal length.

### *In silico* identification of G-quadruplex motif in AST and structural modeling of wildtype and mutant AST sequences

To identify potential G-quadruplex motifs in AST, we used the open-source web-based tool Quadruplex forming G-rich Sequences (QGRS) Mapper (https://bioinformatics.ramapo.edu/QGRS/index.php) (*38*). The search was performed using the following parameters: maximum length 30 nt; minimum tetrads 3 (G_3_N_y1_G_3_N_y2_G_3_N_y3_G_3_); loop size 0-30. The server identified a QGRS of 28 nt at positions 1048-1075 (domain B) of AST with a G-score of 38 (Fig. S5). A second 26-nt QGRS with a G-score of 38 was found that overlaps the first and covers positions 1057-1082 of AST (Fig. S5). The maximum G-score possible is 105 with a G_6_TG_6_TG_6_TG_6_. The 70-nt substitution in ASTmut70B encompasses both sequences. QGRS Mapper did not identify any G-quadruplex motif in ASTmut70B.

### Statistical analyses

Statistical analyses were performed with the Wilcoxon matched pairs signed rank test, and parametric paired and unpaired *t*-tests using Prism GraphPad 10.

---

**Fig. S1.**
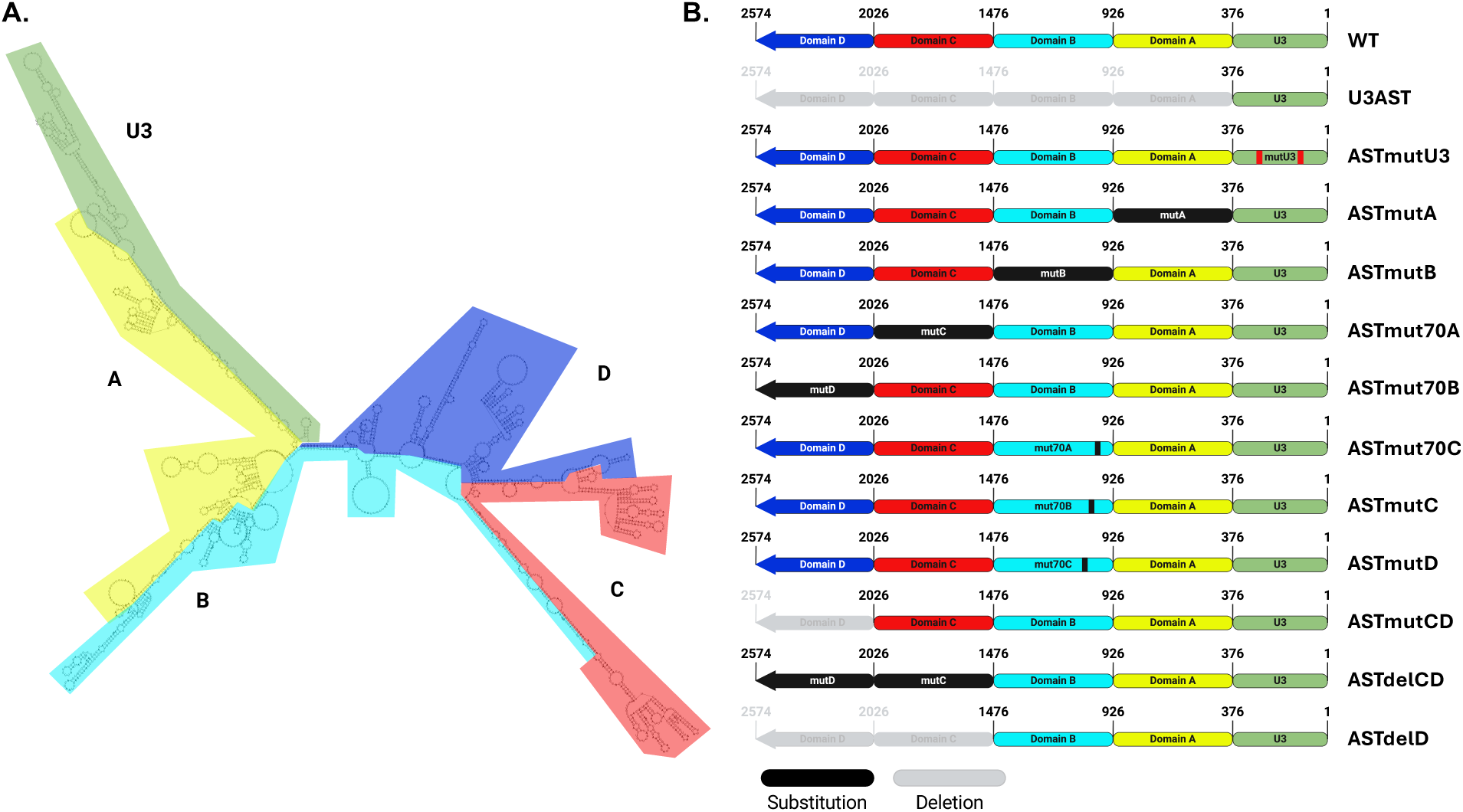
(**A**) Secondary structure (minimum free energy) of wildtype AST predicted with RNAfold (*22–25*). The five domains of AST (U3, A, B, C, and D) are highlighted in green, yellow, teal, red, and blue, respectively. (**B**) Schematic representation of wildtype AST and the panel of AST mutants generated for this study. Domains colored in black indicate substitution with an exogenous sequence of equal length. Domains colored in gray indicate deletions. Substitutions of short motifs in the U3 domain of the ASTmutU3 mutant are indicated in red. Substitutions of short motifs in the B domain of the ASTmut70A, ASTmut70B, and ASTmut70C are indicated in black.

**Fig. S2.**
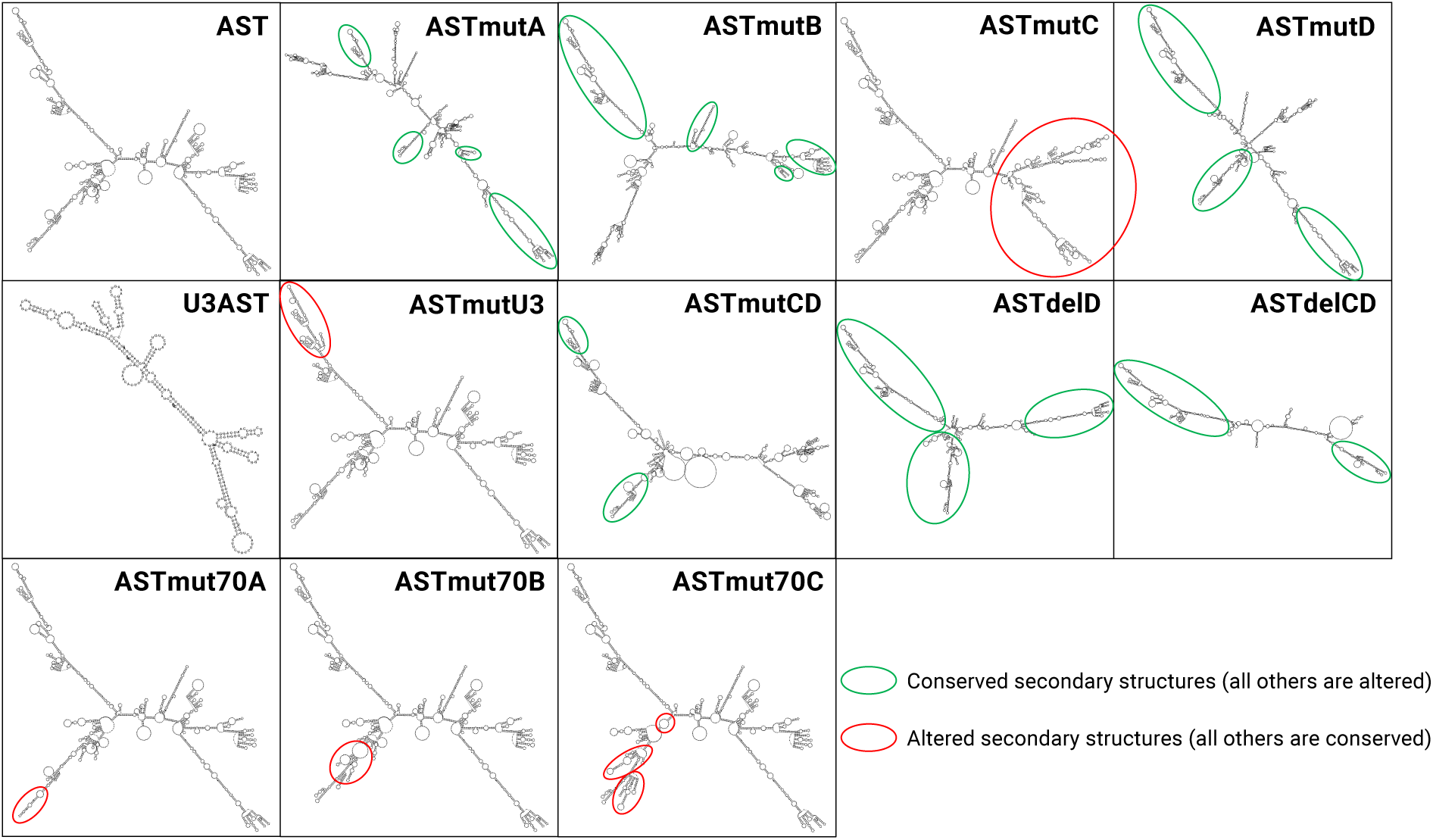
Secondary structure (minimum free energy) of wildtype AST and of the AST mutants predicted with RNAfold (*22–25*). Green circles indicate secondary structures that are conserved in the mutants compared to wildtype AST. Red circles indicate new secondary structures in the mutants compared to wildtype AST.

**Fig. S3.**
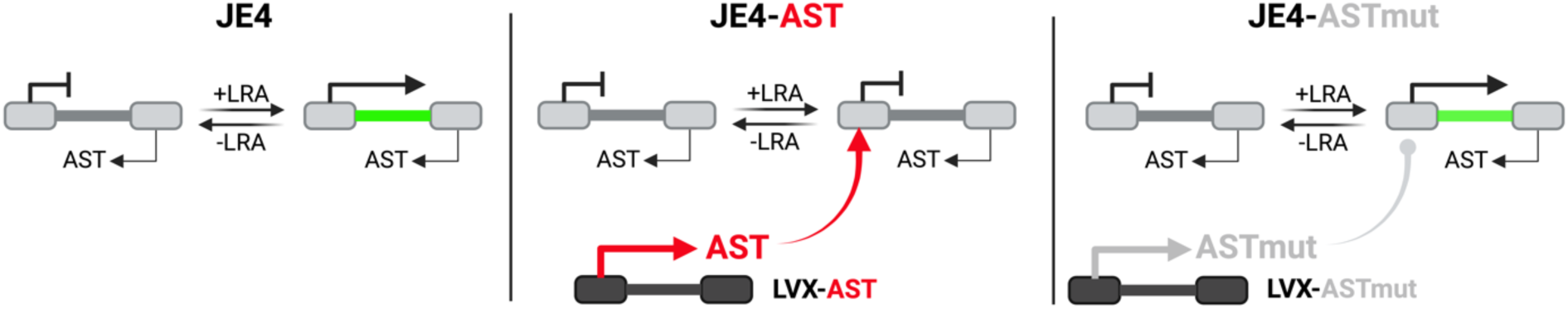
The Jurkat E4 (JE4) latency model. Left: In parental JE4 cells, HIV-1 expression can be triggered with latency reversal agents (LRA) and monitored by GFP expression. Removal of LRA leads to re-establishment of latency. The HIV-1 provirus naturally expresses low levels of AST (*19*). Middle: JE4 cells stably transduced with a lentiviral vector expressing high levels of wildtype AST (JE4-AST). This cell line is used to study the effects of AST on HIV-1 expression (*19*). Right: JE4 cells stably transduced with a lentiviral vector expressing high levels of different AST mutants (JE4-ASTmut).

**Fig. S4.**
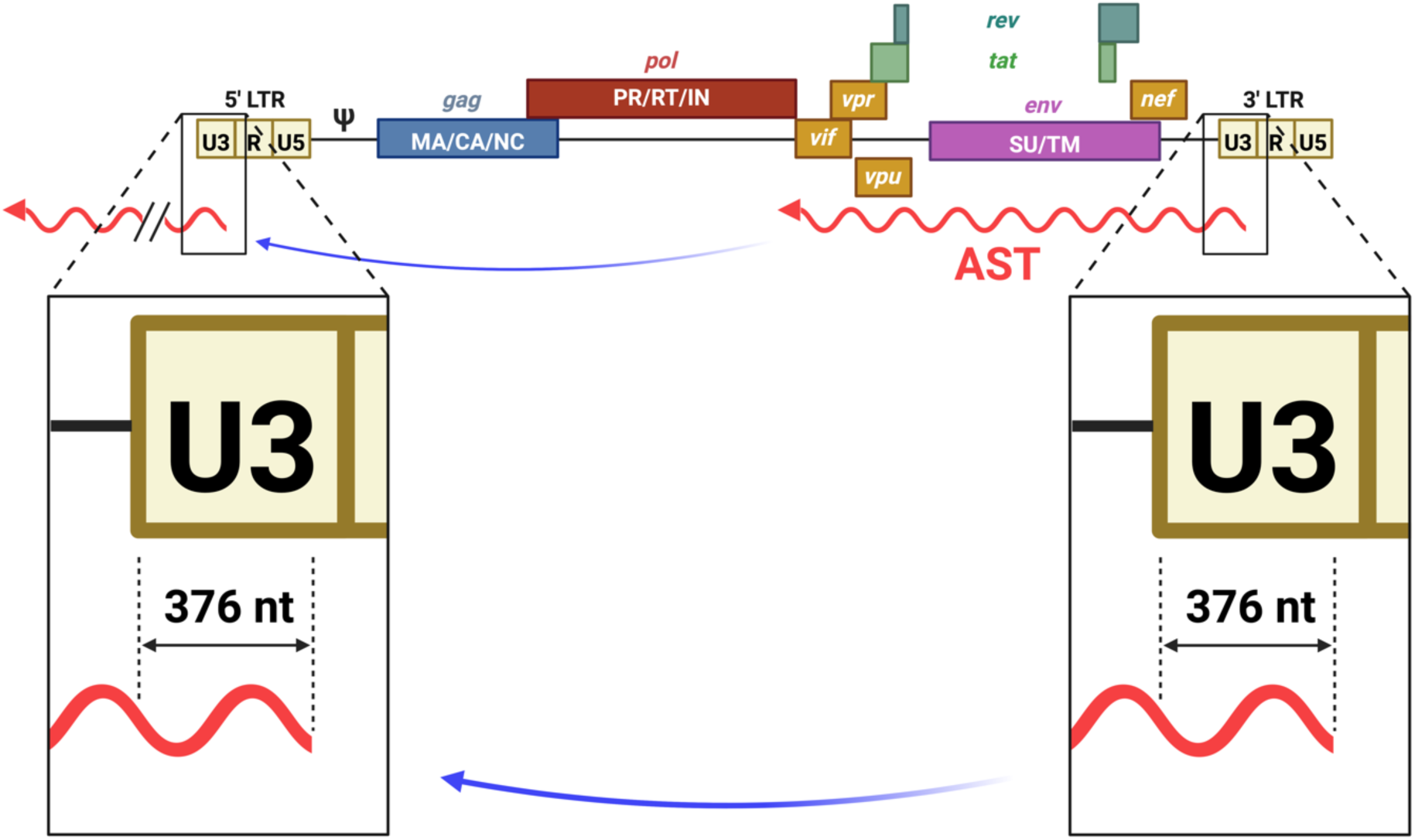
Diagram of the HIV-1 proviral genome and the antisense transcript AST. The 5’ terminus of AST covers 376 nt of the U3 region in the 3’ LTR (zoom-in inset on the right) and it has perfect sequence homology with the U3 region in the 5’ LTR (zoom-in inset on the left).

**Fig. S5.**
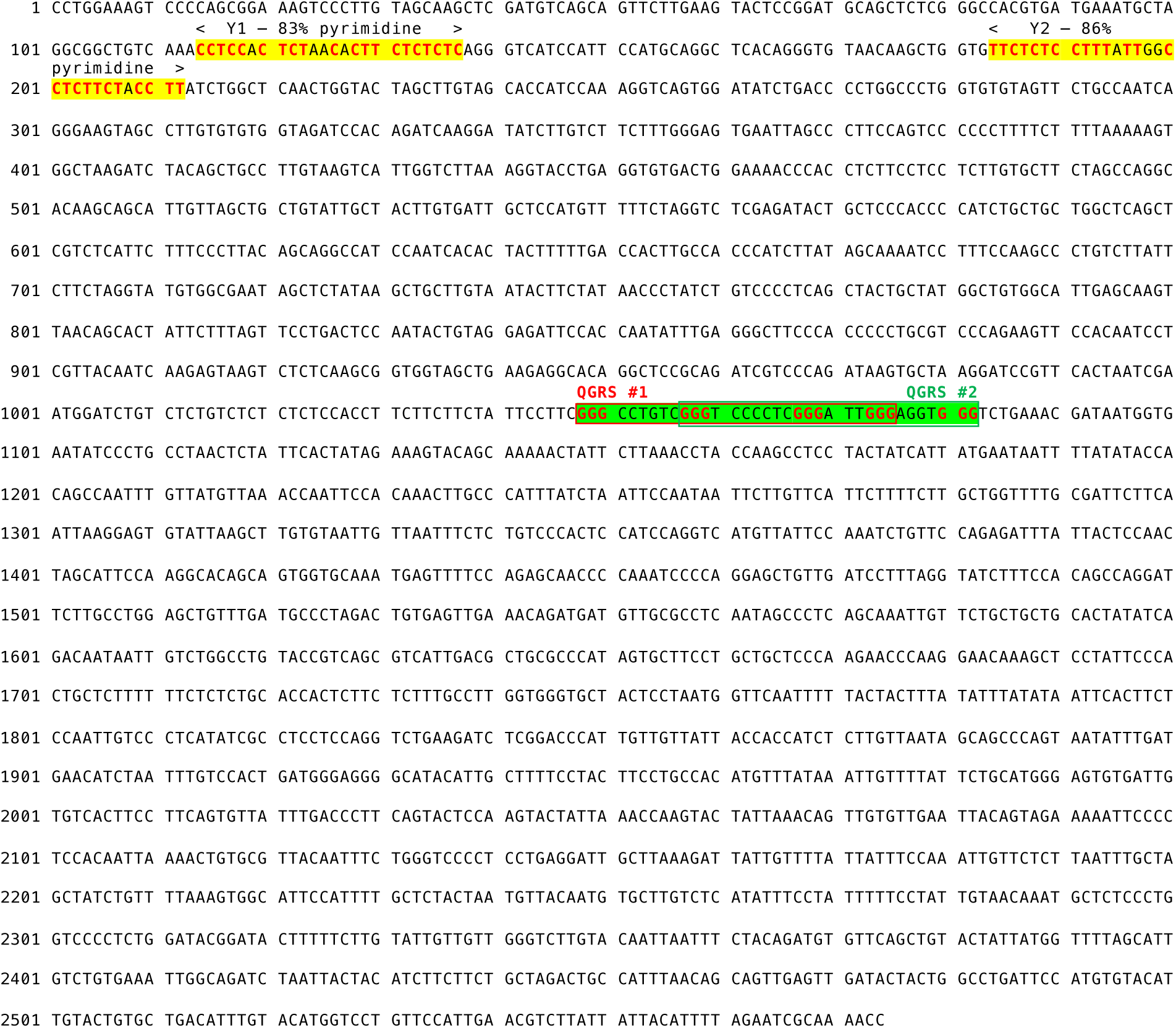
Position of critical functional motifs withing the sequence of wildtype AST. Motifs highlighted in yellow indicate the two pyrimidine-rich Y1 and Y2 motifs in the 5’ terminus of AST (U3 domain). Pyrimidine residues are shown in red, bold letters. The sequence highlighted in green indicates two overlapping QGRS motifs in domain B of AST: motif #1 is enclosed in red-lined box and motif #2 is enclosed in green-lined box. G-triplets are shown in red, bold letters.

**Fig. S6.**
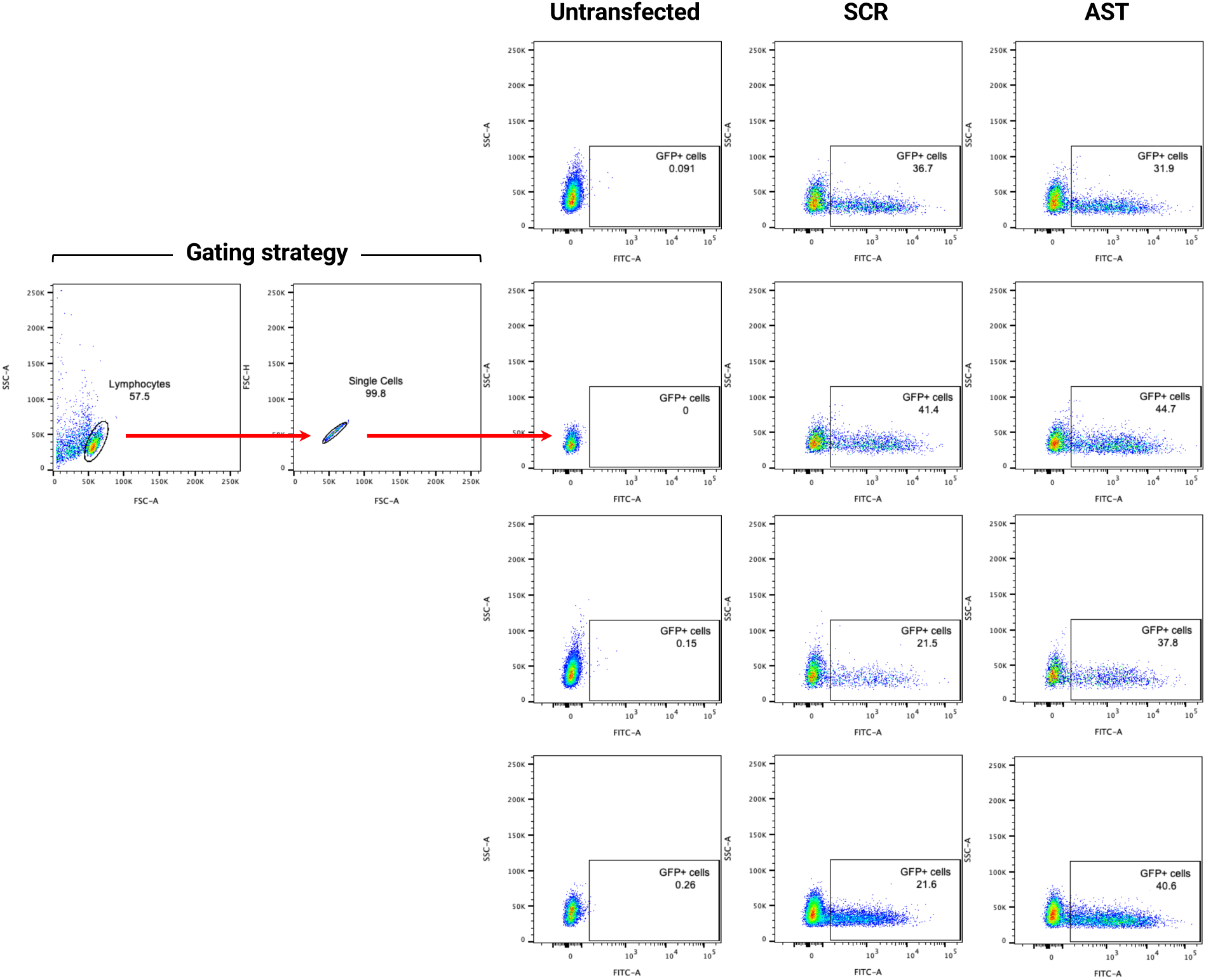
Efficiency of nucleofection by GFP expression. Flow cytometry analyses of GFP expression in CD4+ T cells from PLWH either untransfected (four plots in the left column) or transfected with SCR-GFP (four plots in the middle column) or AST-GFP vectors (four plots in the right column). The two plots at the far left show the gating strategy that was used to assess the percentage of GFP+ cells (FITC-A).

**Fig. S7.**
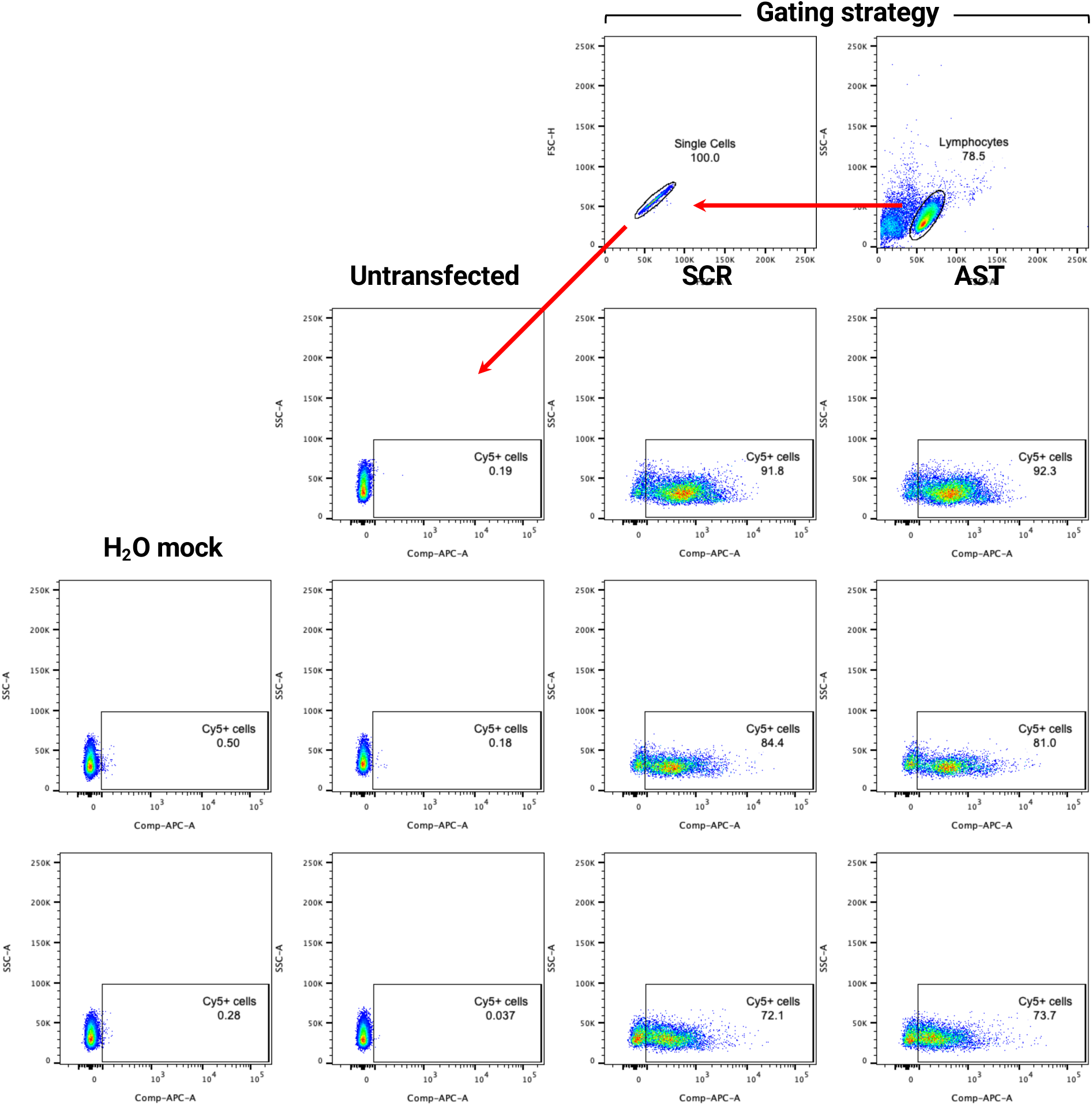
Efficiency of nucleofection by DNA-bound Cy5 uptake. Flow cytometry analyses of CD4+ T cells from PLWH either untransfected (three plots in the second column) or transfected with either SCR-GFP (three plots in the third column) or AST-GFP vectors (three plots in the fourth column) after labeling with Cy5. As a control for uptake of residual Cy5 dye not bound to plasmid DNA, sterile water was “mock labeled” with Cy5, and then used to transfect CD4+ T cells (two plots in the first column). The two plots at the top show the gating strategy that was used to assess the percentage of GFP+ cells labeled with Cy5 dye.

**Table S1.**
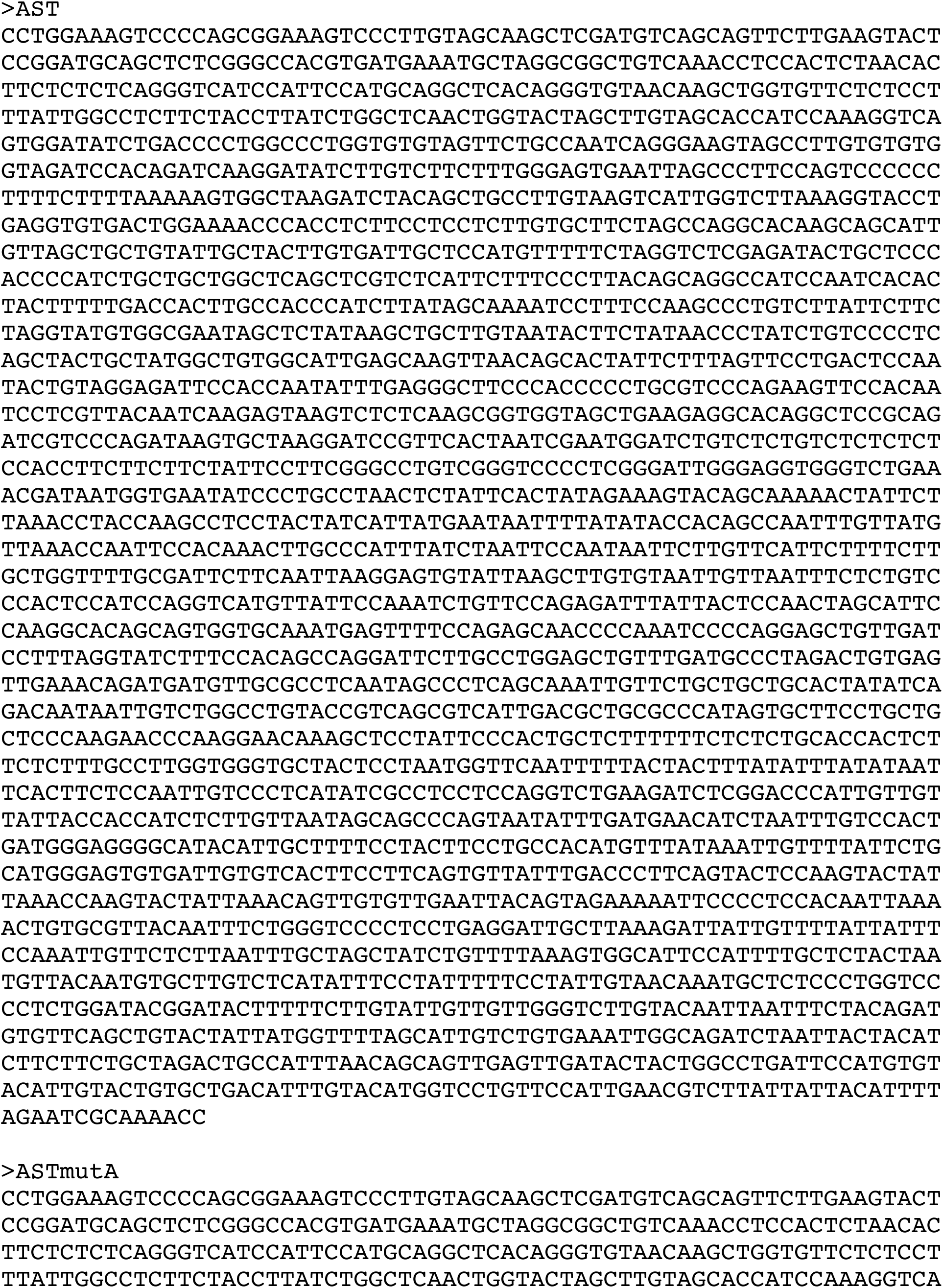
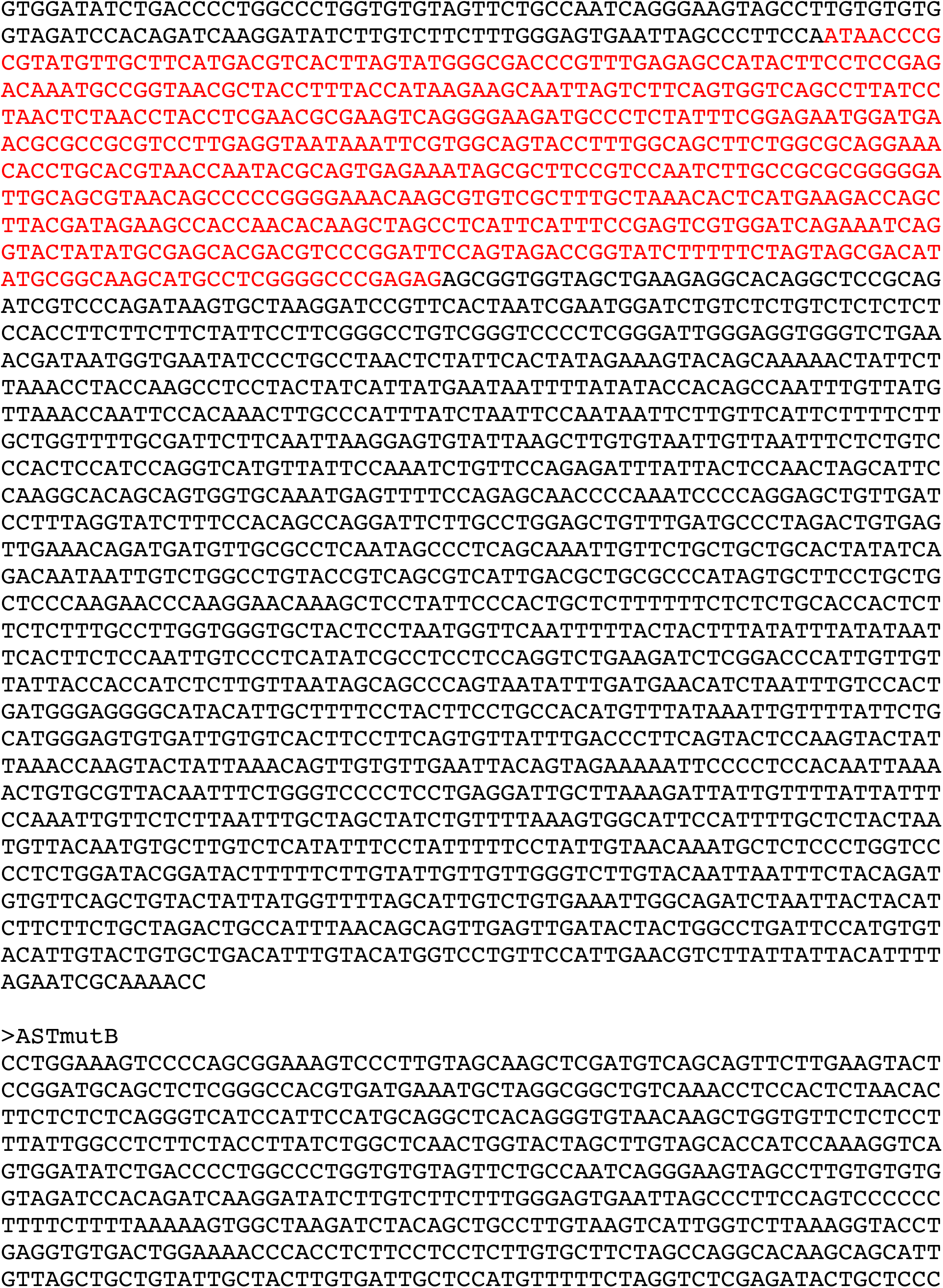
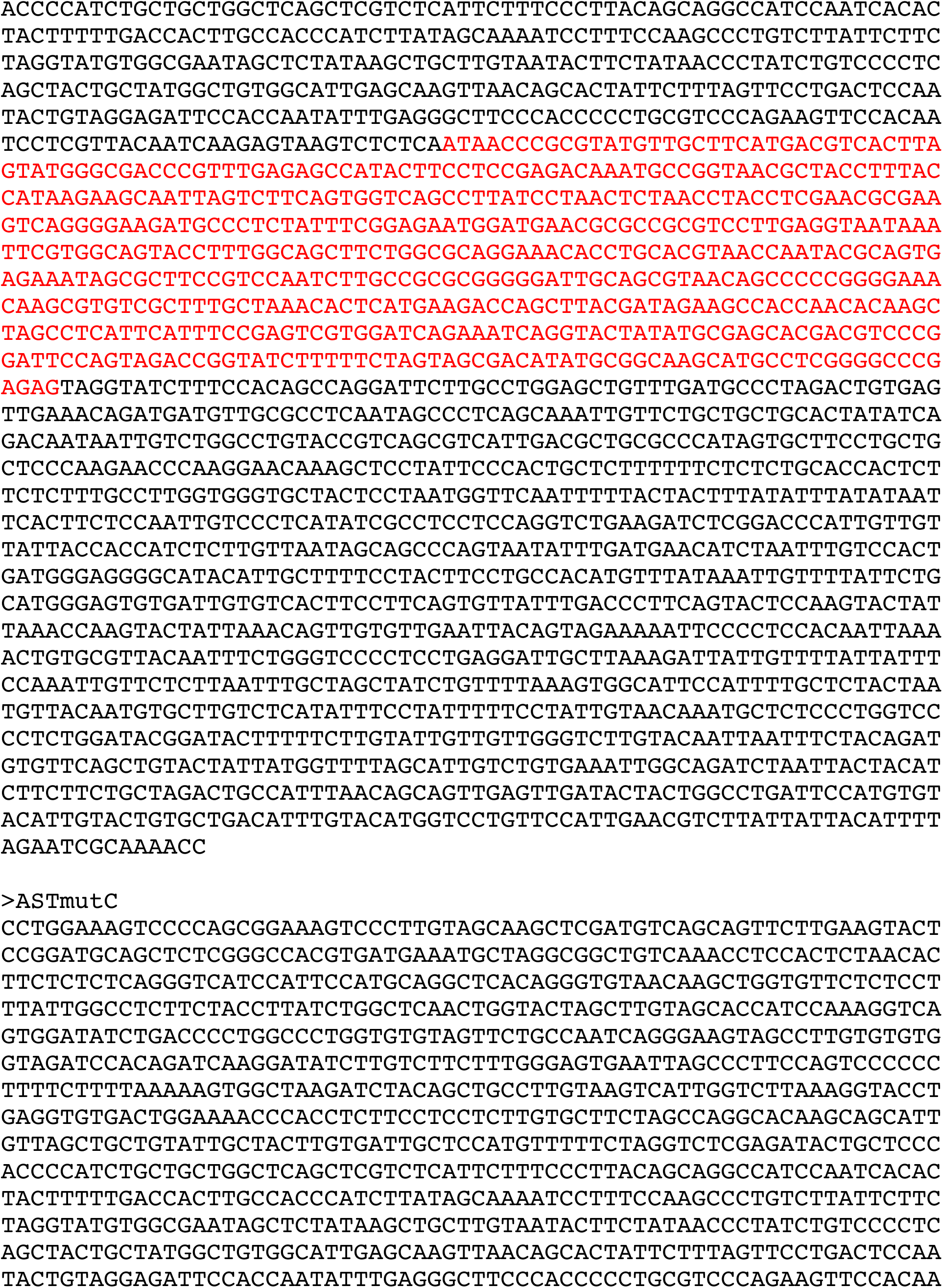
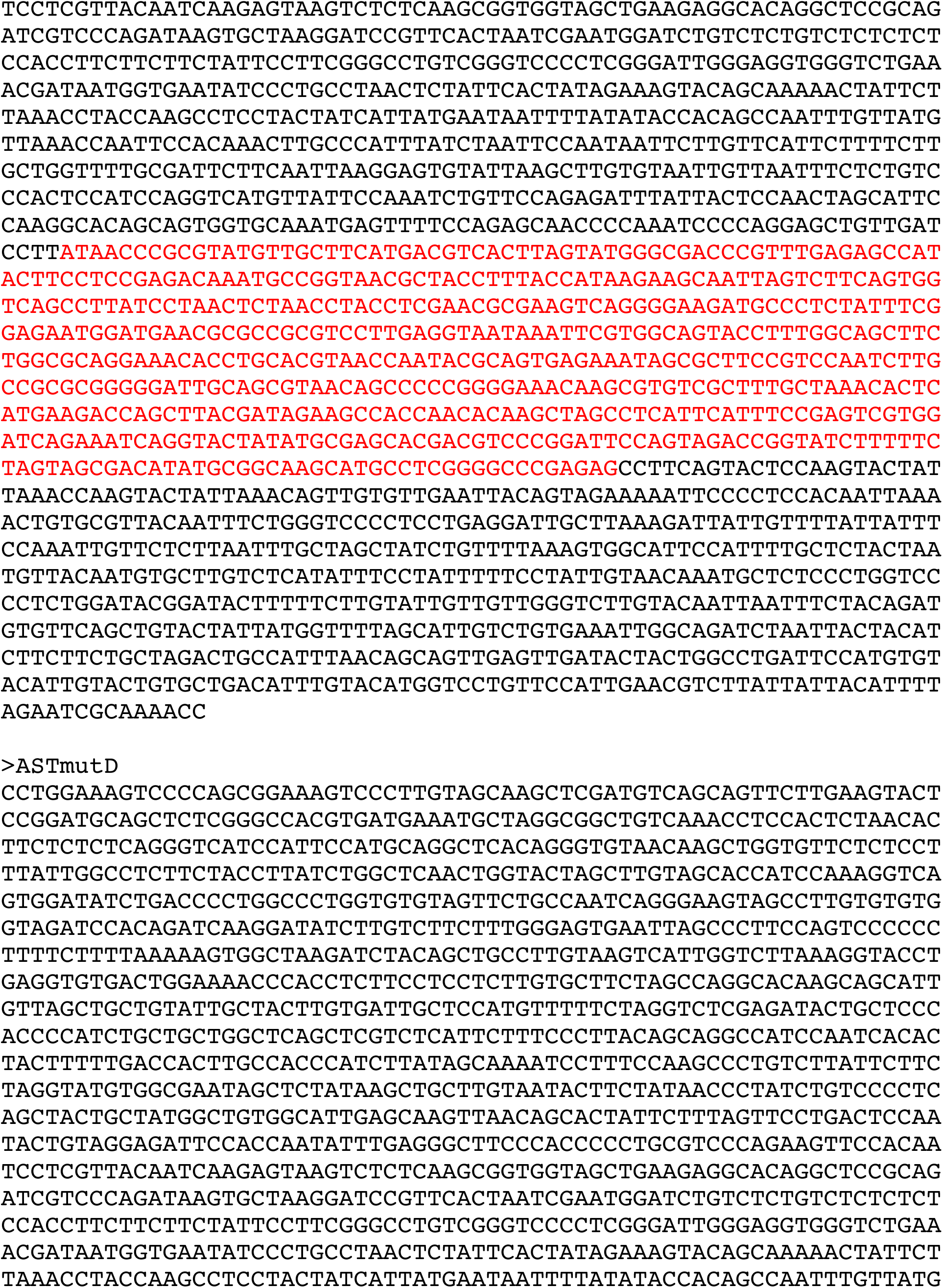
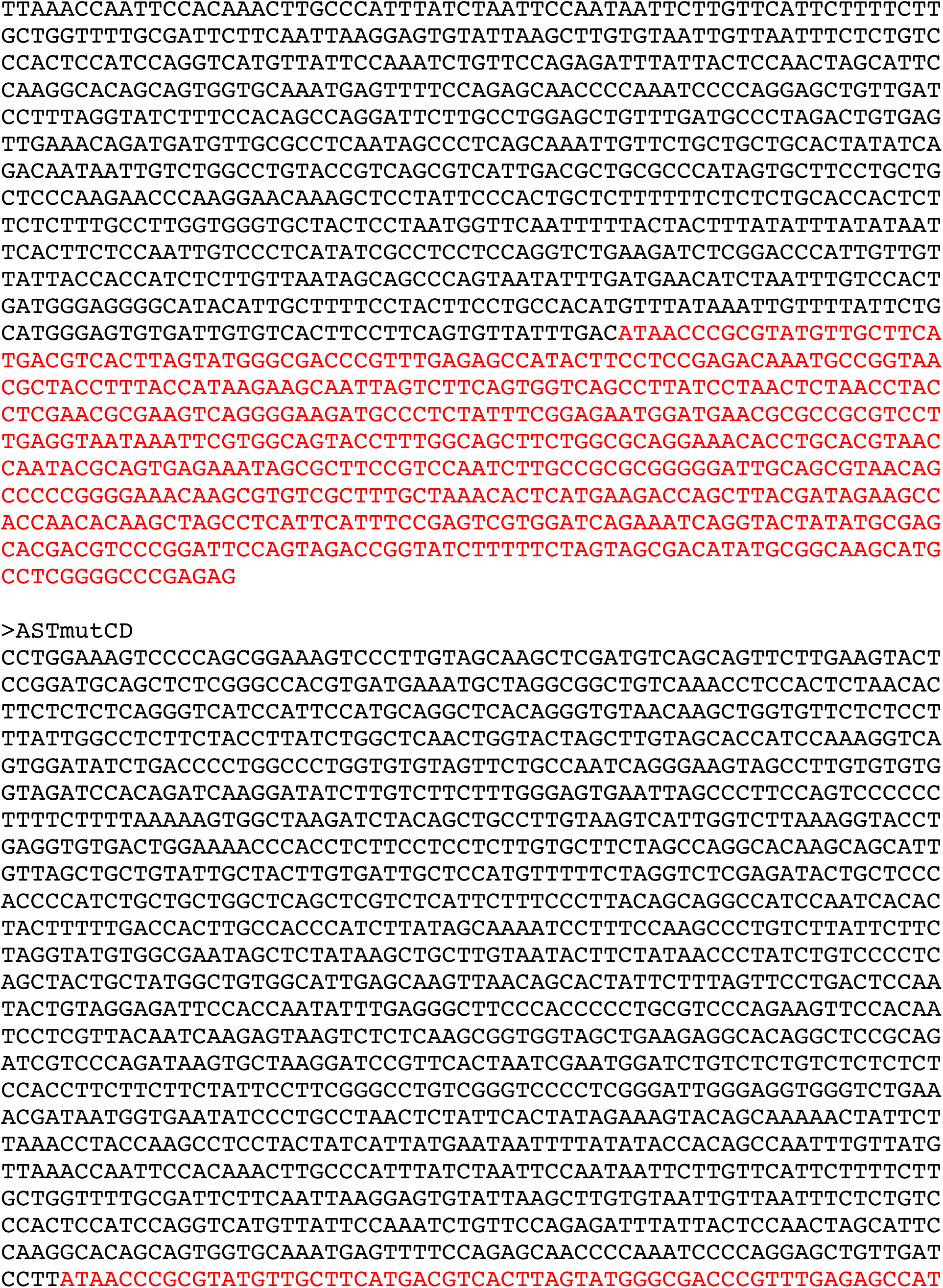
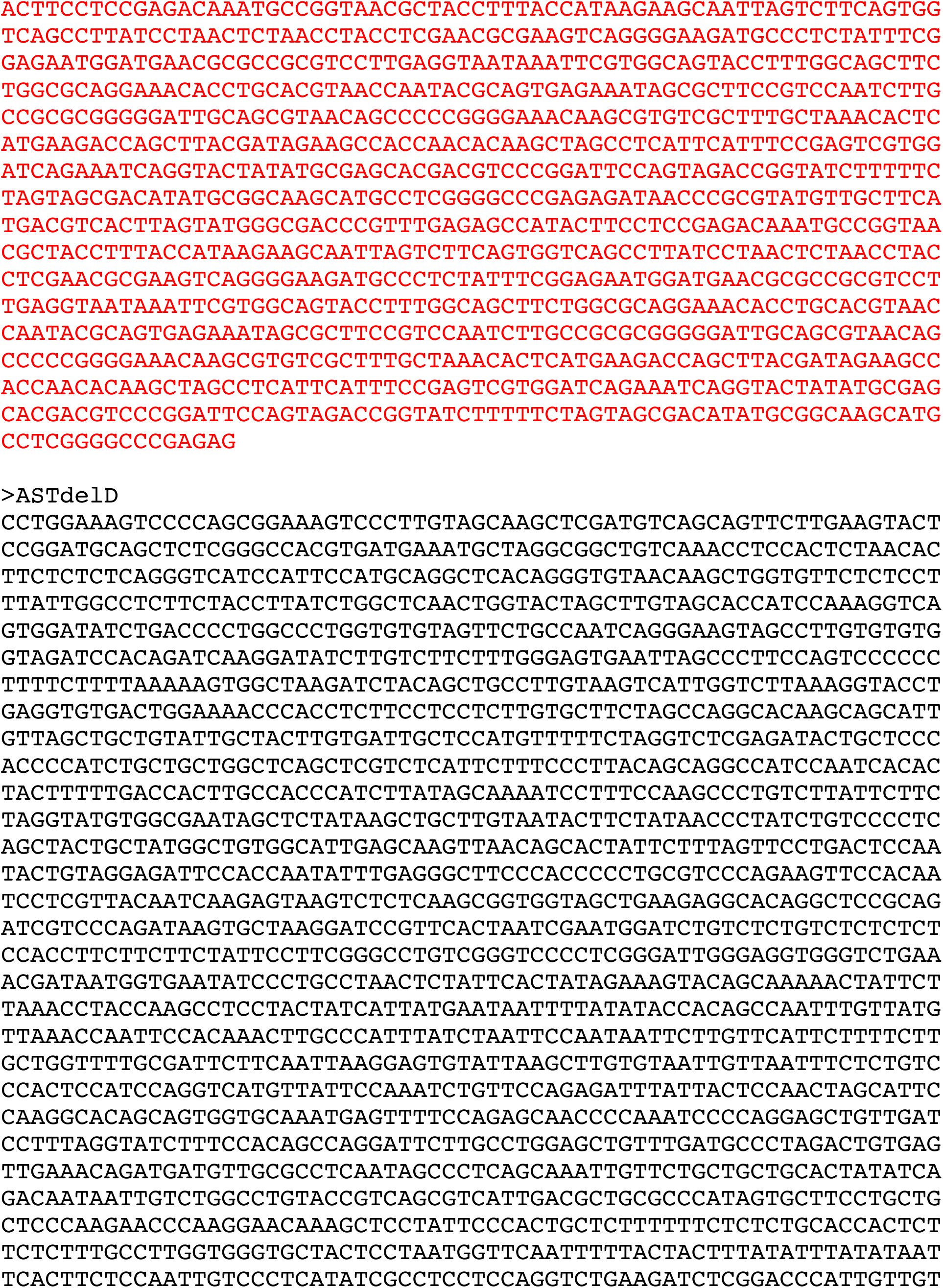
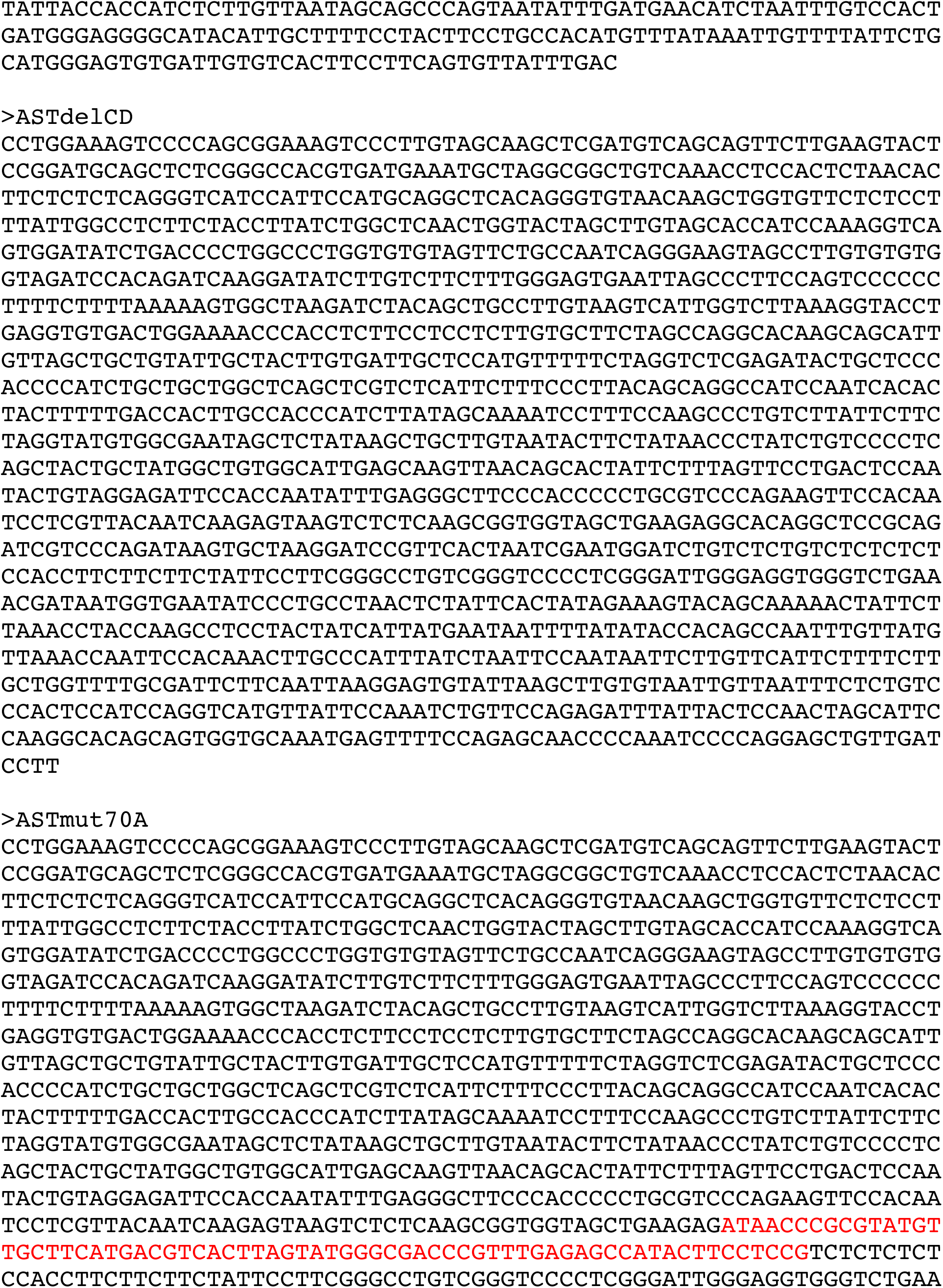
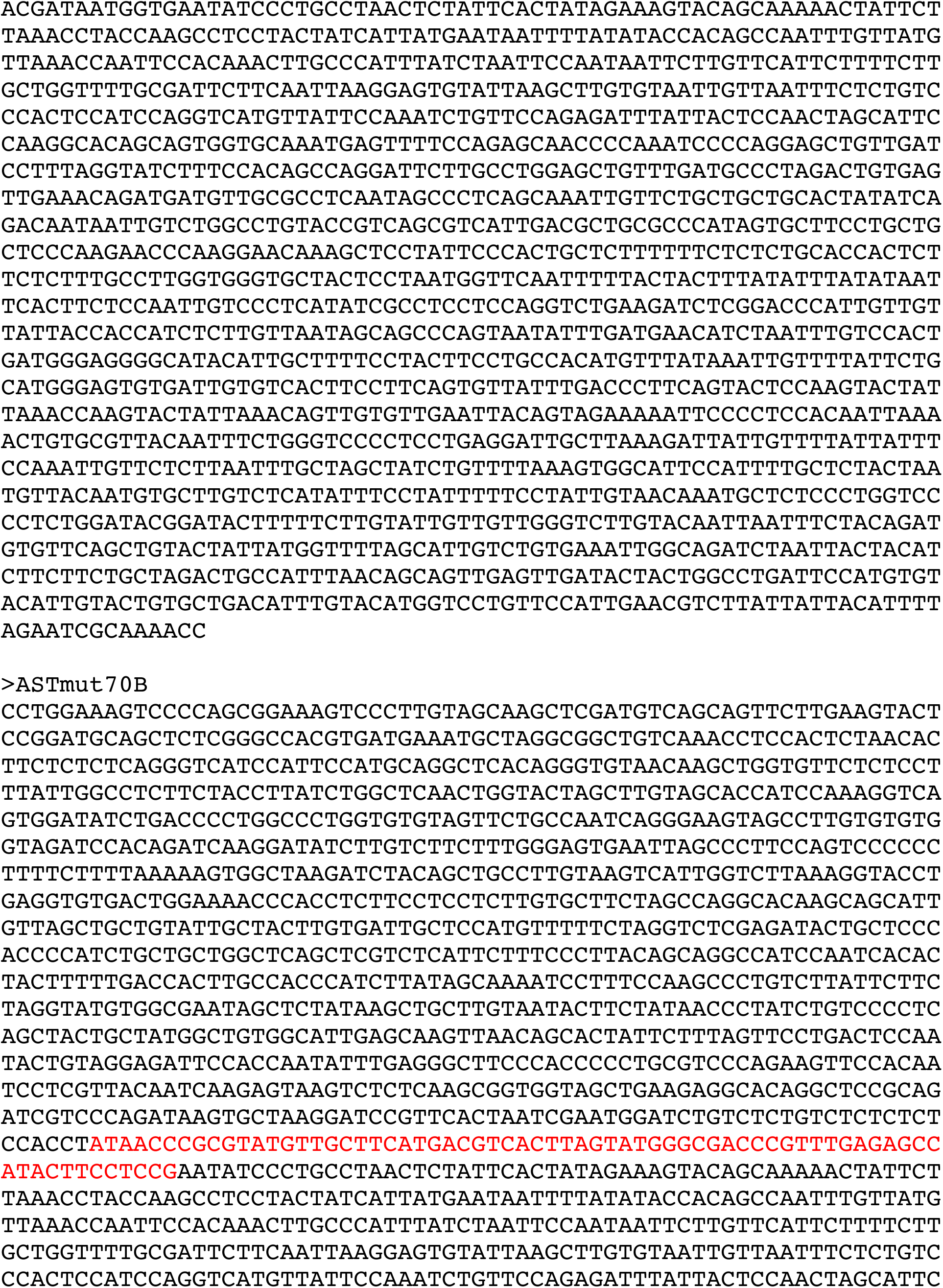
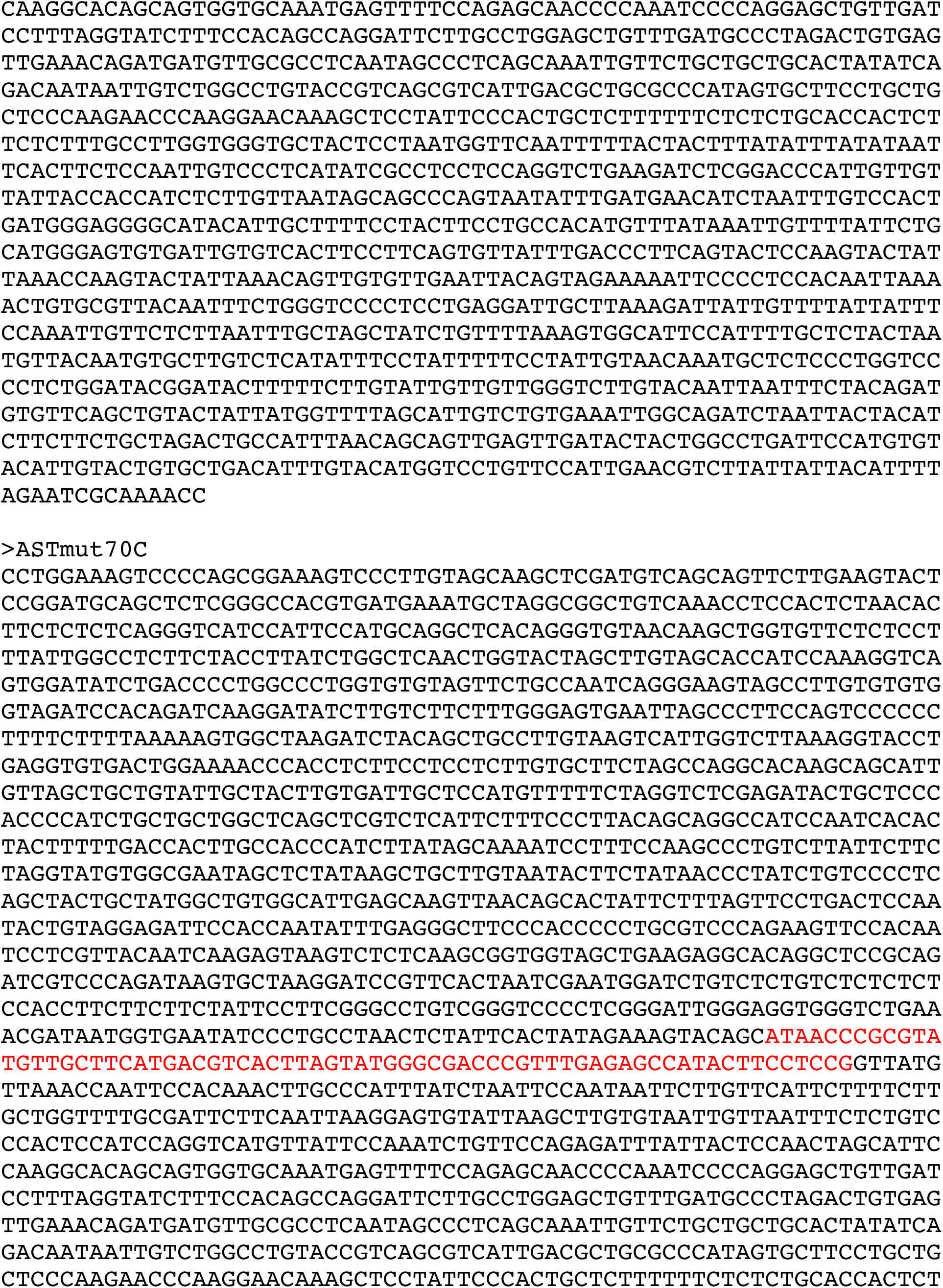
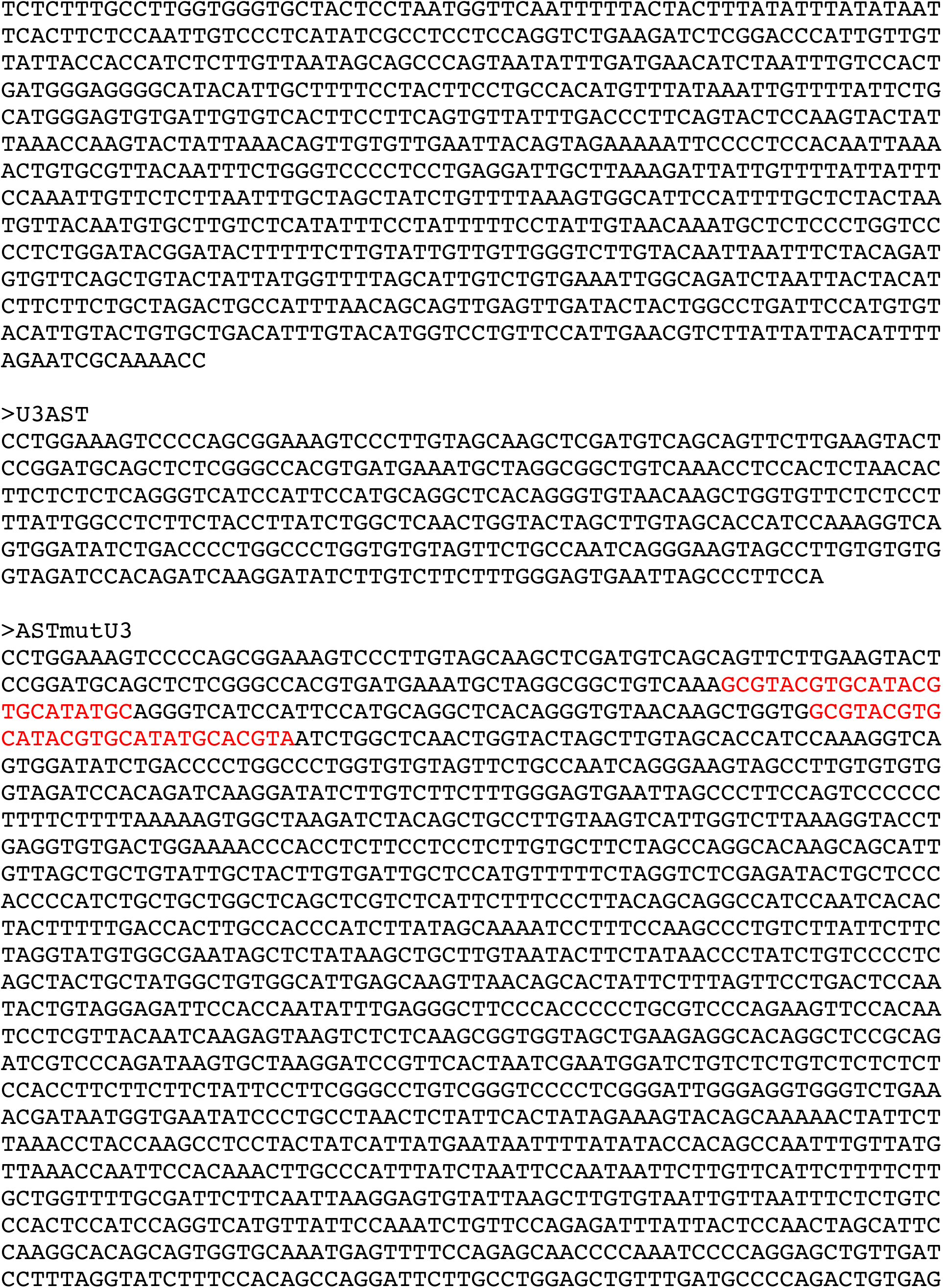
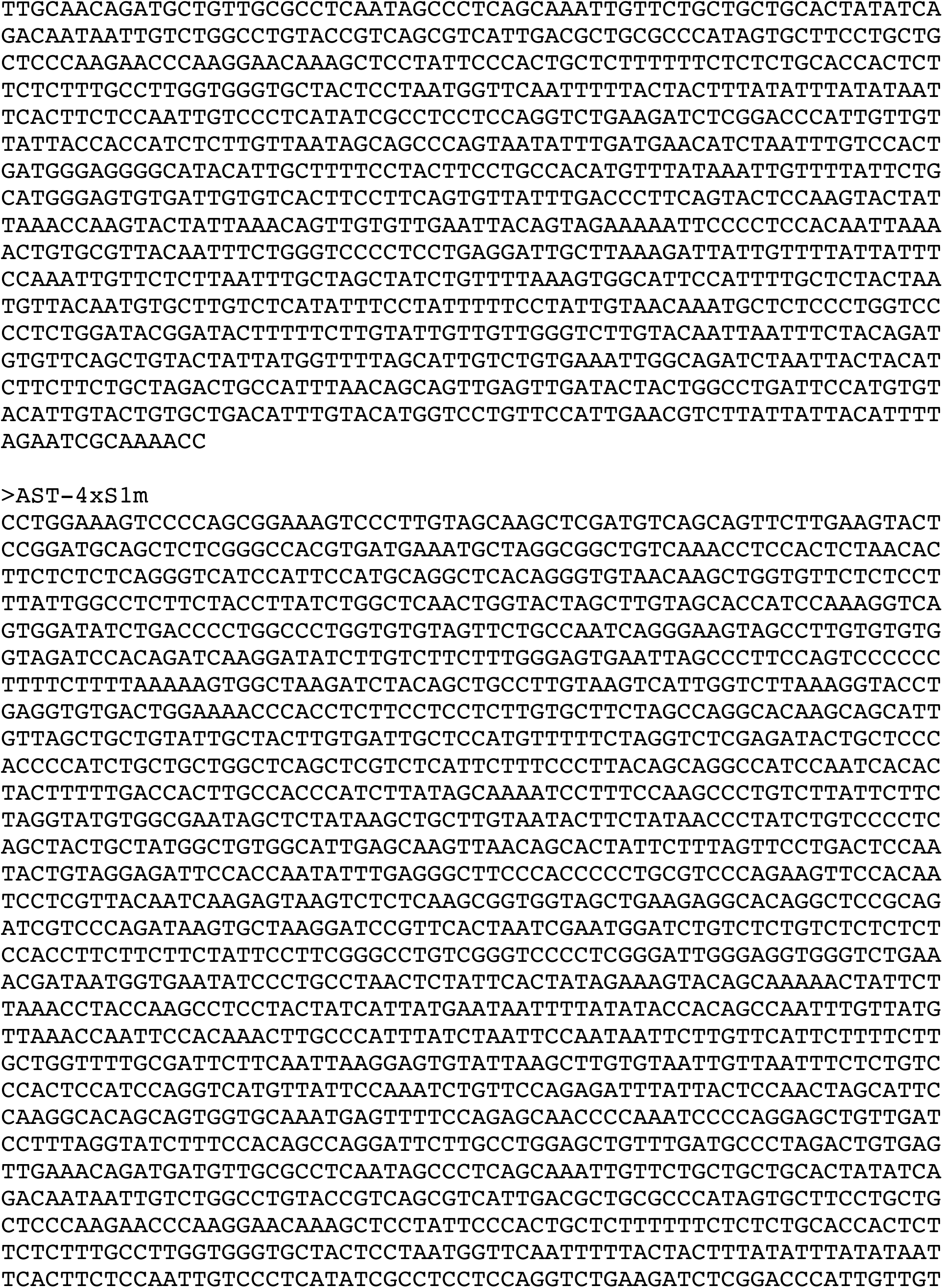
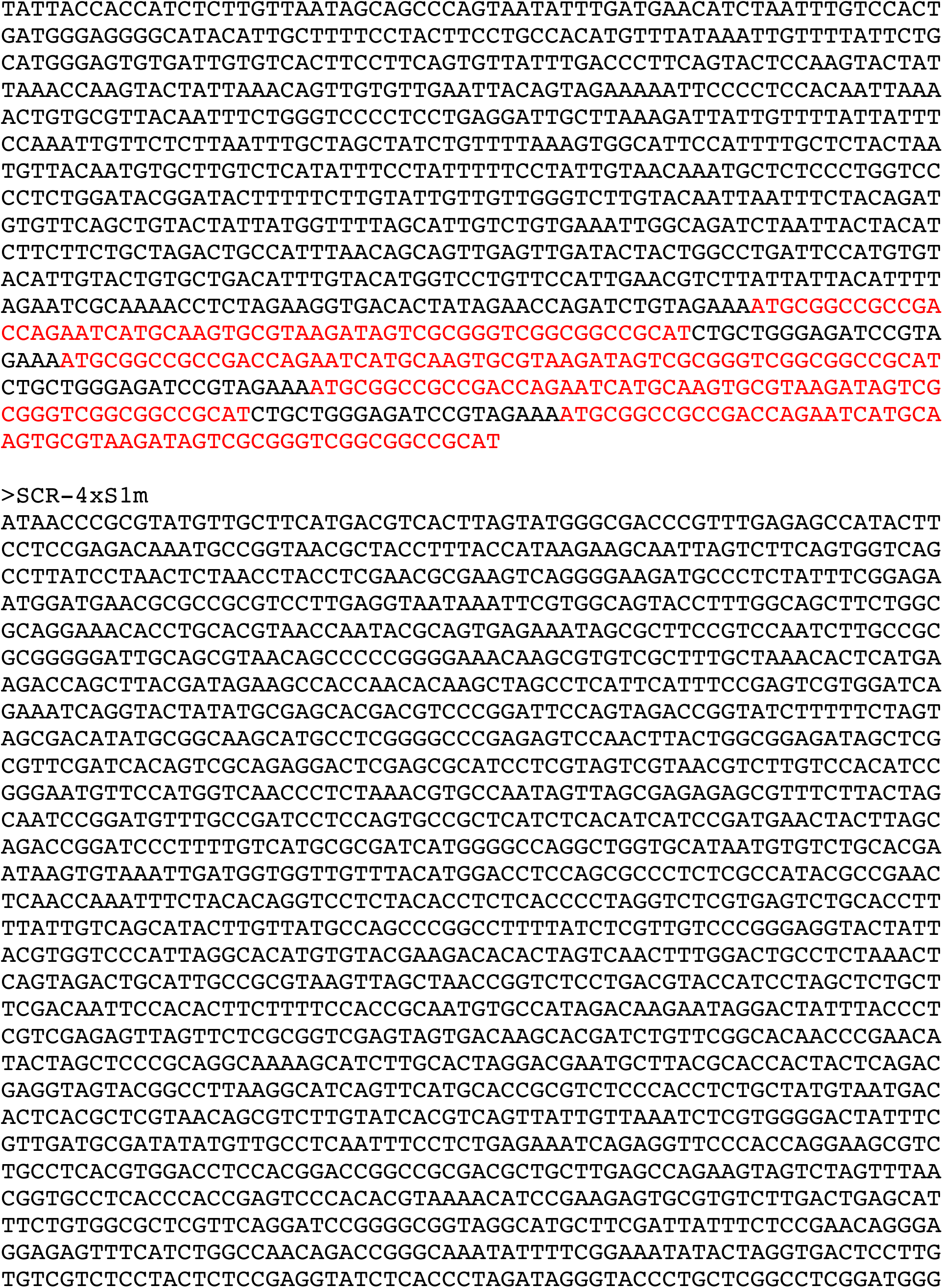
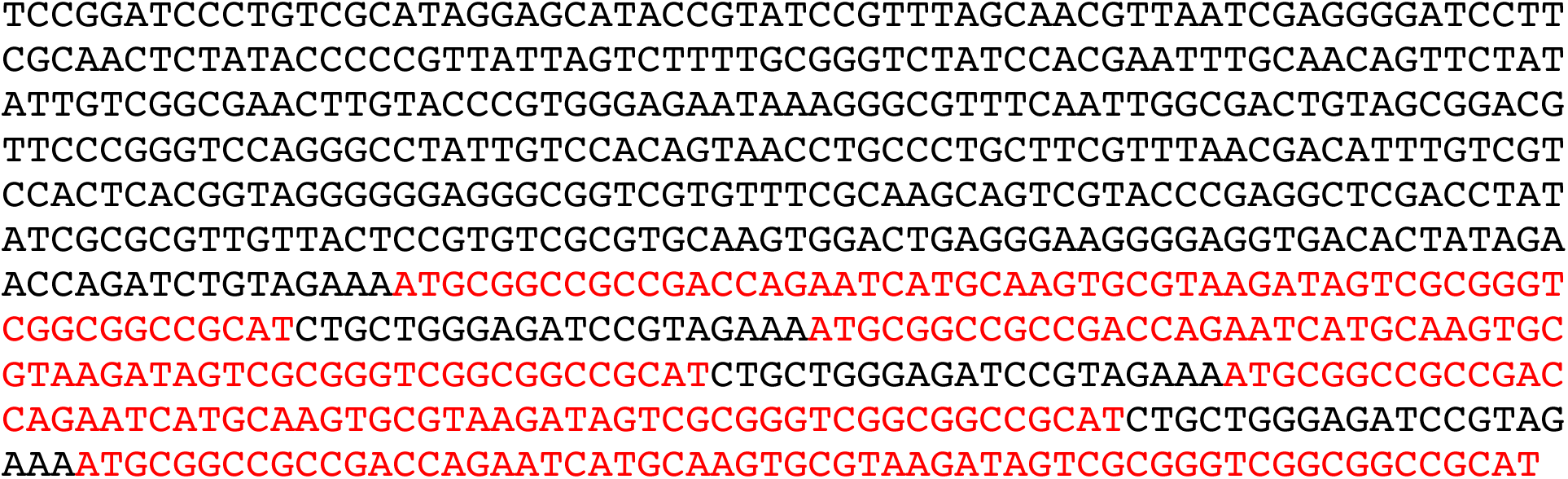
Sequence of wildtype AST and AST mutants utilized in this study. All mutants were generated by DNA synthesis (GeneScript), cloned into the lentiviral vector pLVX-Puro (Takara) under the EIF1α promoter, stably transduced into JE4 cells, and maintained under puromycin selection (*13*). Sequences highlighted in red indicate the exogenous sequence that was used to replace domains or motifs in the wildtype AST. The same exogenous sequence was used to generate the ASTmutA, ASTmutB, ASTmutC, and ASTmutD mutants. In the ASTmutCD mutant, two copies of the exogenous sequence were placed in tandem after domain B in place of domains C and D. To generate the ASTmut70A, ASTmut70B, and ASTmut70C mutants, we used the first 70 nt of the exogenous sequence inserted in the mutants above. The ASTmutU3 mutant was generated by replacing the Y1 (residues 114-137; 83% pyrimidine content) and Y2 motifs (residues 184-212; 86% pyrimidine content) with sequences of alternating purine-pyrimidine residues (50% and 48% pyrimidine content, respectively) of equal length. The sequences highlighted in red in the AST-4xS1m and SCR-4xS1m sequences indicate the four S1m aptamers.

**Table S2.**
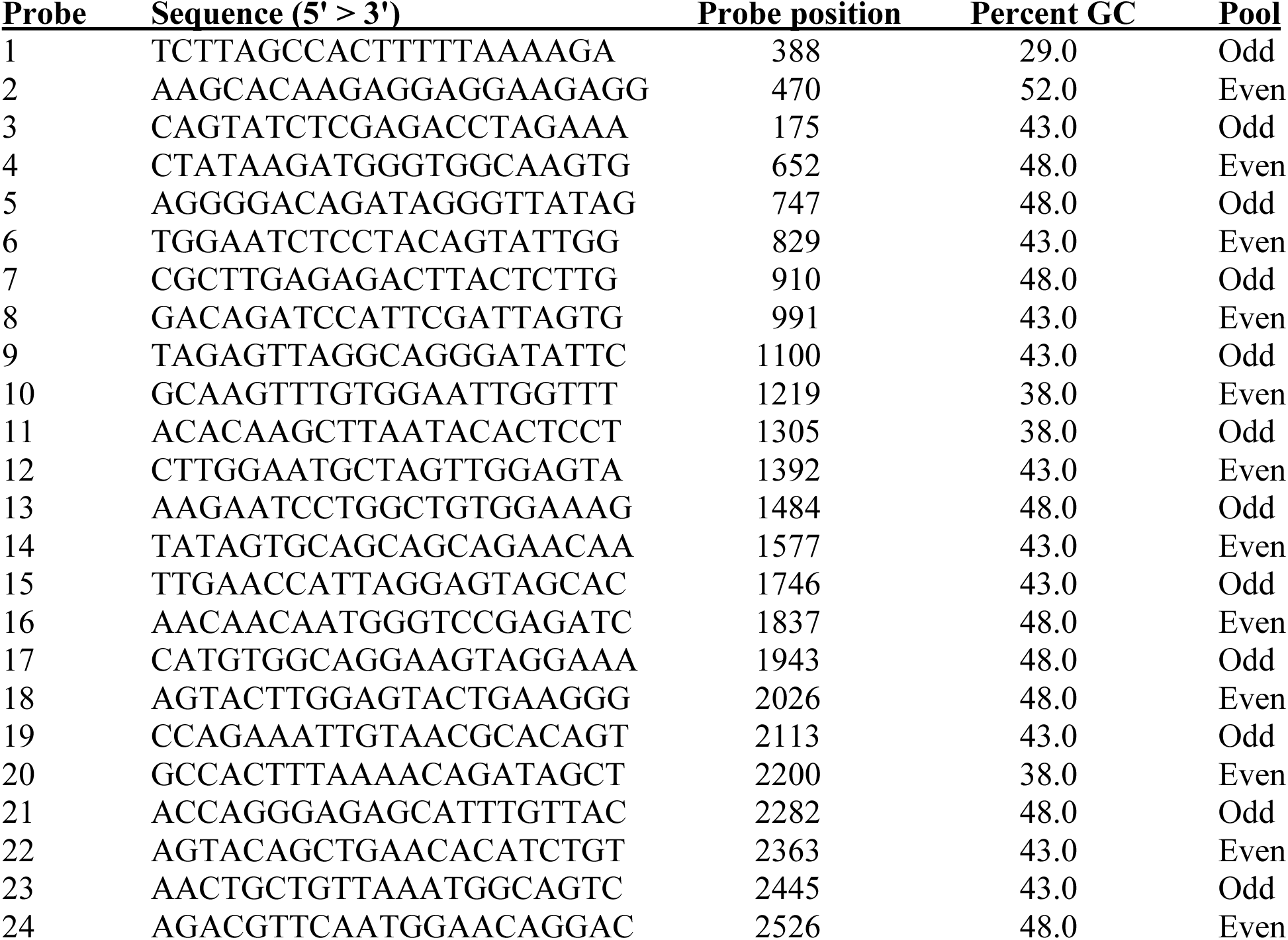
Oligonucleotide probes used for ChIRP. Probes were designed on the portion of the AST sequence between position 377 and 2574, which excludes the U3-derived sequence containing two polypyrimidine-rich stretches. Probes were generated using the online design tool at www.biosearchtech.com. The table shows the 5’ > 3’ sequence of each probe (reverse complementary of the AST sequence), the position of the probe on the AST sequence (the last residue in the probe sequence pairs with the indicated position on the AST sequence), and the probe pool (Odd or Even) that each probe was assigned to.

**Table S3.** Donors demographics.

| **Participant ID** | **Age (Years)** | **Sex** | **Race** | **ART Regimen** | **Years of continuous suppression at time of sampling** |
| --- | --- | --- | --- | --- | --- |
| FR38 | 71 | M | Black | DTG, RPV | 1 |
| FR43 | 65 | M | Black | BIC, FTC, TFV | 2 |
| FR51 | 57 | M | Black | BIC, FTC, TFV | 1 |
| FR52 | 51 | F | Black | DRV, COBI, FTC, TFV | 5 |
| FR53 | 68 | M | Black | DTG, 3TC | 6 |
| FR54 | 55 | M | Black | BIC, FTC, TFV | 9 |
| FR55 | 69 | M | Black | ABC, DTG, 3TC | 10 |
| FR65 | 66 | F | Black | DTG | 5 |
| FR67 | 33 | F | Black | CAB, RPV | 13 |
| FR70 | 63 | M | Black | DRV, TFC, TFV, RTV, DTG | 6 |
| FR72 | 37 | F | White | CAB, RPV | 3 |
| FR73 | 60 | M | Black | EVG, CAB, FTC, TFV | 18 |
| FR74 | 38 | M | Black | CAB, RPV | 7 |
| FR77 | 44 | M | Black | BIC, FTC, TFV | 7 |
| FR78 | 44 | F | Black | BIC, FTC, TFV | 2 |

**Table S4.**
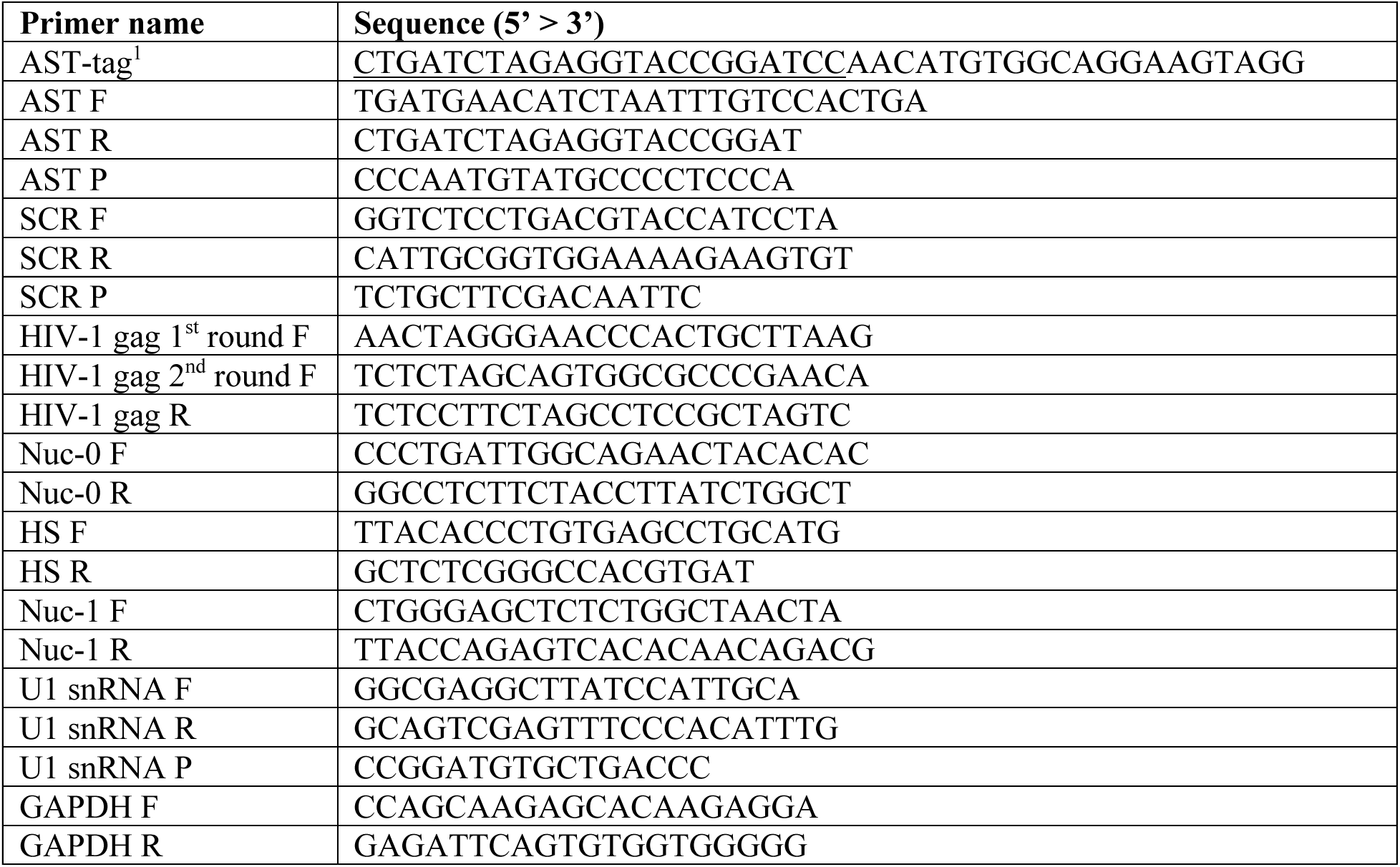
Sequence of primers and probes for RT and qPCR. F: forward; R: reverse; P: probe. ^1^Underlined sequence identifies the exogenous tag in the RT primer

**Table S5.** Antibodies used for RIP and ChIP assays.

| **Target** | **Host** | **Clone** | **Supplier** | **Cat. #** |
| --- | --- | --- | --- | --- |
| Total IgG (for RIP) | Rabbit | Polyclonal | Millipore | PP64B |
| TDP-43 | Rabbit | Polyclonal | Proteintech | 10782-2-AP |
| EZH2 | Rabbit | EPR20108 | Abcam | AB191250 |
| hnRNPK | Rabbit | Polyclonal | Bethyl | A300-674A |
| Matrin 3 | Rabbit | Polyclonal | Bethyl | A300-591A |
| PTBP1 | Rabbit | E4I3Q | Cell Signaling Technology | 57246S |
| SMARCD2 (Baf60B) | Rabbit | Polyclonal | Millipore | ABE1318 |
| Total IgG (for ChIP) | Rabbit | C2C12 | Millipore | 05-623-25UG |
| RNA Polymerase II | Mouse | CTD4H8 | Cell Signaling Technology | 4620S |
| H3K27me3 | Rabbit | C36B11 | Cell Signaling Technology | 9733S |

**Data S1.**
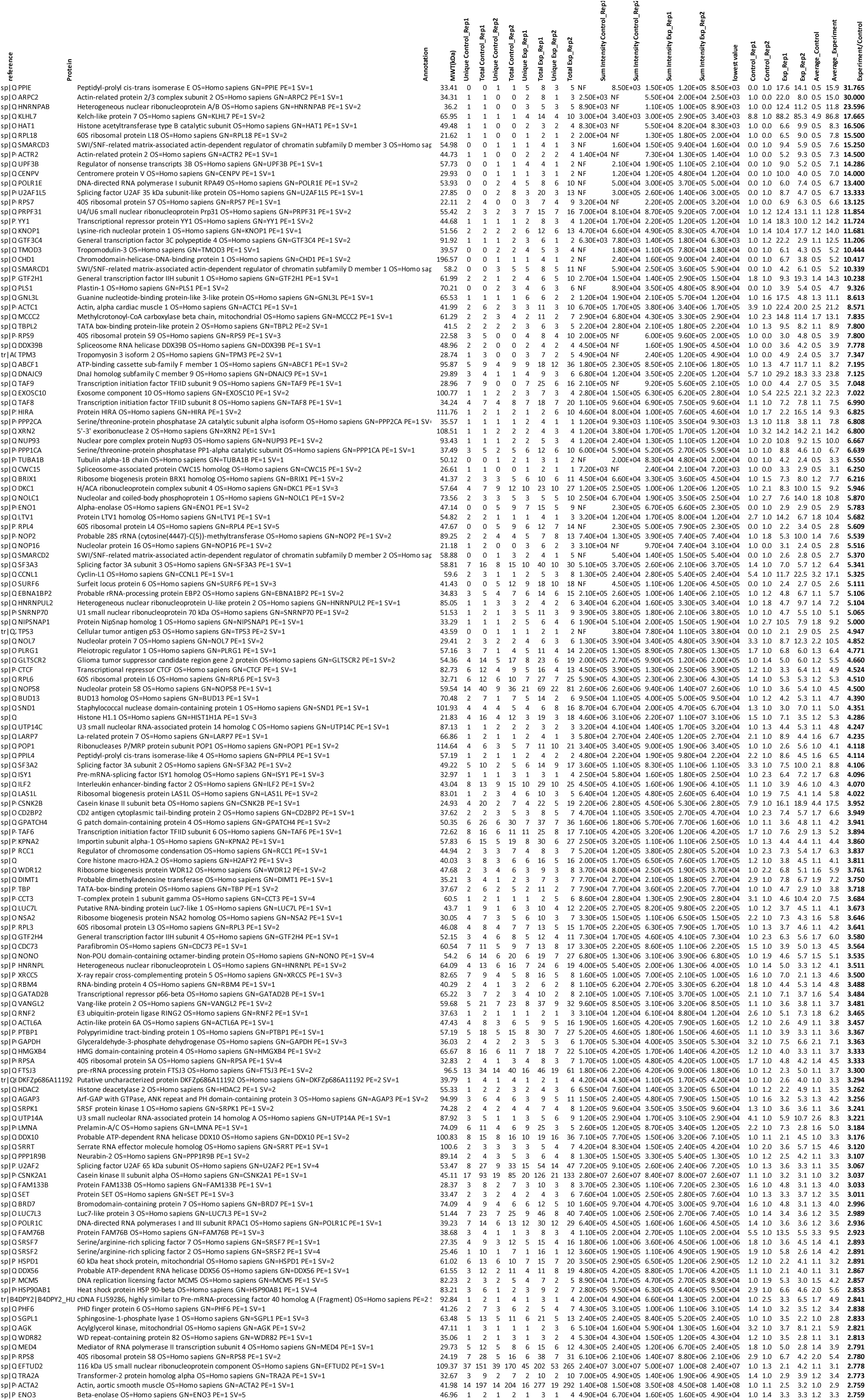
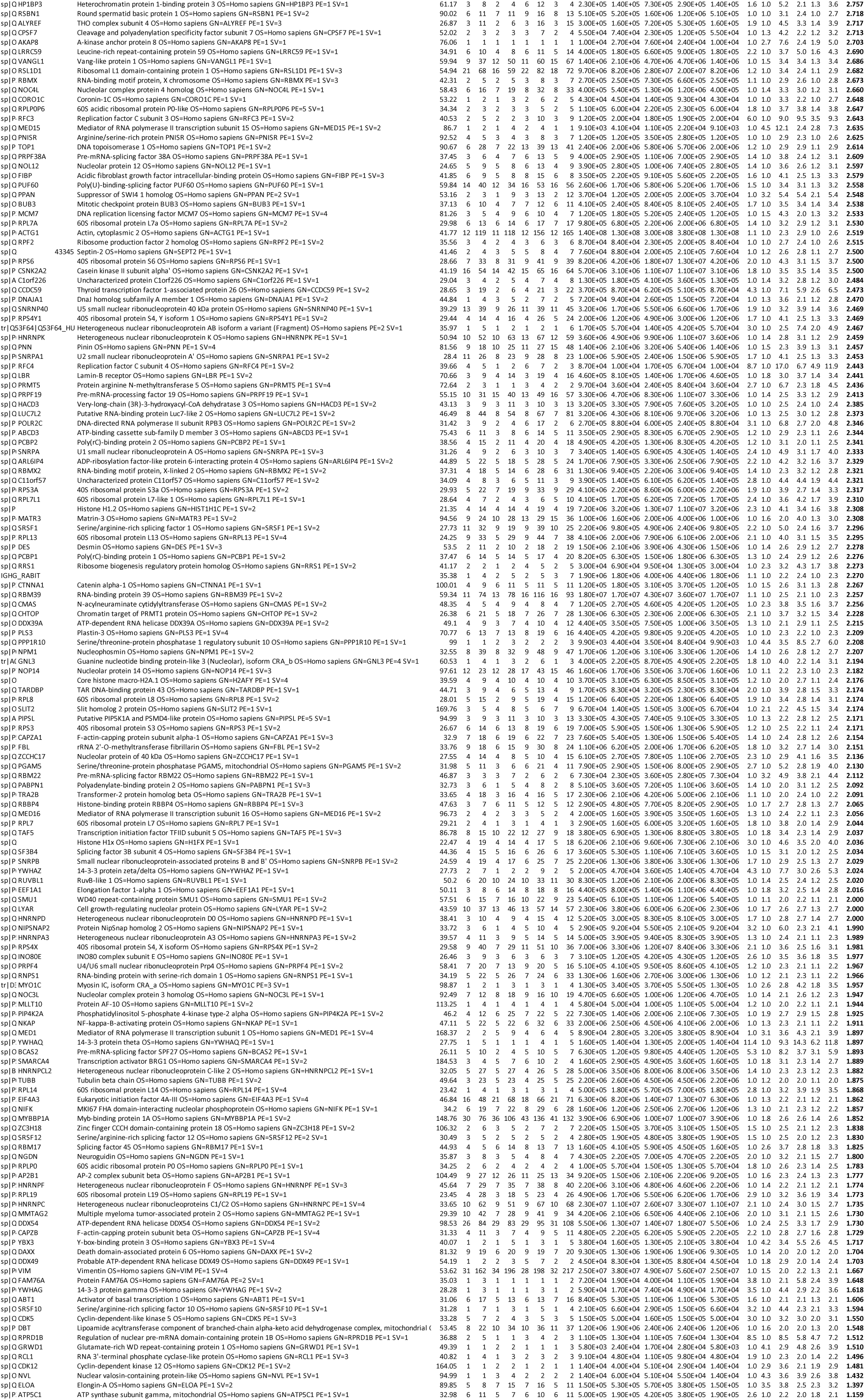
Excel spreadsheet listing host proteins enriched in the AST-4×S1m pulldown compared to the SCR-4×S1m pulldown.

